# Functional randomness despite high taxonomic turnover across an elevational gradient in a global biodiversity hotspot: A case study of hawkmoths and birds

**DOI:** 10.1101/867770

**Authors:** Mansi Mungee, Ramana Athreya

**Author notes:** **Author contributions:** MM formulated the hypotheses, performed the analyses and drafted the manuscript. RA is the principal investigator of the larger on-going biodiversity project in the region. The collection and curation of field and trait data, were shared equally by both. Wildlife Institute of India, P.O.Box # 18; Chandrabani Dehrdun 248001, India.

## Abstract

**Aim:** We examined the patterns and processes of taxonomic and functional dissimilarities for two disparate organismal groups (ectothermic hawkmoths and endothermic birds) across a broad tropical elevational gradient.

**Location:** Eaglenest Wildlife Sanctuary (northeast India), eastern Himalayan global biodiversity hotspot.

**Taxon:** 4,731 hawkmoths; 15,387 birds

**Methods:** Turnover and nestedness components for taxonomic and functional dissimilarities were obtained using the methods developed by Baselga (2013) and Leprieur et al., 2012. We used Generalized Dissimilarity Modeling (GDM) with geographic distance, contemporary and historic climatic variables to assess the relative importance of dispersal and environmental processes in determining the beta diversity. Functional redundancy (FRed) was calculated for both organismal groups using the Simpson’s diversity indices. Null modeling was used to determine randomness in species and trait distributions.

**Results:** Turnover dominated taxonomic and functional dissimilarities, however the contribution of nestedness was considerably higher to the latter. Overall, the rate of dissimilarity with distance, for both facets of diversity, was significantly higher for birds, with stronger contributions of geographic distance and historic climate; whereas the hawkmoth dissimilarities were strongly correlated with only contemporary climate. Taxonomic dissimilarities deviated significantly from null, whereas functional dissimilarities exhibited high redundancy and randomness.

**Main Conclusions:** Overall, our results suggest that while the drivers of beta-diversity exhibit idiosyncrasy and taxon-specificity; for a given taxa, they are consistent across the two facets of dissimilarity. More importantly, regardless of the principal predictor, the net result was that of high taxonomic turnover, which is de-coupled to a high degree from functional turnover in these tropical ecosystems. The large redundancy in trait values, despite high species turnover, indicates functional resilience of these tropical communities. The consistency of this pattern, across two disparate organismal groups, is suggestive of a key mechanism in which tropical communities may retain functionality of ecosystems in a changing environment.

## Introduction

Beta diversity, or the compositional difference among communities, is a central concept in ecology and has received a renewed interest due to its pivotal role as a link between local (alpha) and regional (gamma) diversity (Buckley & Jetz, 2008). Especially within the context of tropical forests, where complete inventories on even local richness are seldom available for a wide variety of organisms, an understanding of the mechanisms generating the spatio-temporal variation in community composition, i.e. beta diversity, has provided valuable insights into the understanding of the mechanisms maintaining the high biodiversity in these regions (KÖnig et al., 2017). A common approach for investigating patterns in beta diversity across environmental gradients is by characterizing the Distance-Dissimilarity-Relationships (hereafter DDRs) i.e., the slope of the relationship between compositional (taxonomic/functional) dissimilarities and environmental/geographic distances (Nekola & White, 1999). Difference in slopes for different taxa has been explained by contrasting dispersal abilities, vagililty and environmental tolerance/niche width (all three inversely related to the slope of DDRs; Soininen et al., 2007). Central to the analysis of DDRs are four major conceptual and methodological developments, each concerning itself with assessing the relative role of a mechanism from a pair of non-exclusive (complementary or antagonistic) processes responsible for generating and maintaining patterns in beta diversity.

The first, and perhaps the most well documented, methodological advancement relates to partitioning the total observed dissimilarity into it’s two integral components – turnover and nestedness (Baselga, 2010, 2013). Compositional dissimilarity between any pair of communities can arise either due to turnover i.e. species replacement, or nestedness i.e. species loss. Different assembly processes are responsible for either type of change (environmental filtering versus extinction-colonization dynamics) and thus partitioning beta diversity into these two components can further the understanding of ecological drivers of dissimilarity (Soininen et al., 2018).

The second methodological development concerning DDRs deals with a more pervasive statistical challenge – spatial auto-correlation of environmental variables. Community dissimilarity can arise either due to dispersal limitation (taxon-specific dispersal) or due to environmental filtering (selection imposed by abiotic conditions, taxon specific niche-width), and consequently disentangling the relative contribution of environment and geographic distances (Tilman, 1982; Hubbell, 2001) can provide important insights into patterns of beta diversity. However, since most environmental variables are strongly correlated with geographic distances, the relative influence of these two filters remains largely unresolved (Qian & Ricklefs, 2007).

Globally, the Quaternary climatic changes have been shown to shape the current patterns of species distributions and diversity across a broad range of organisms (Araújo et al., 2008, Hortal et al., 2011, Svenning et al., 2015), but it’s relative contribution, in comparison with contemporary climate and spatially limited dispersal, especially across the Himalayas has been rarely investigated (Yang et al., 2008, Yu et al., 2015). The Himalayan glaciation was affected considerably during the Late Quaternary period (Owen, Derbyshire & Fort, 1998, Owen, Finkel & Caffee, 2002). Geological evidences and Global Climatic Models (GCMs) reveal glaciation till up to 10 km further from contemporary ice boundaries during the Last Glacial Maximum (LGM), which reduced the monsoon precipitation in southeast Asia, and affected the distribution of many taxa (Owen et al., 2002). Thus, disentangling the relative contribution of historic and contemporary climate to observed patterns in composition is critical for predicting the fate of biodiversity in the light of emerging climate changes. The recent advancements in the assembly of historic climatologies have paved the way for this third important analytical advancement in the understanding of mechanisms generating beta diversity (Gent et al., 2011; Watanabe et al., 2011; Fitzpatrick et al., 2013; Giorgetta et al., 2013).

Finally, ecological communities respond to the changing environment in not just the number, type and abundance of the constituent species, but also in their functional trait composition (Lamana et al., 2014). Taxonomic and functional dissimilarities are expected to be positively correlated due to the principles of limiting similarity, which predict a minimum permissible overlap across the niche space of two co-occurring species (MacArthur & Levins 1967). A redundancy in trait composition despite high taxonomic turnover may indicate ecosystem resilience to perturbations and environmental changes (Swenson et al., 2011). Similarly, a disproportionately higher loss in functional diversity, in comparison to taxonomic diversity may make communities more vulnerable to climate change (Robroek et al., 2017). Especially useful in this context have been the examinations of deviations between the expected and observed dissimilarities using null modeling approaches (Díaz et al., 2007; Cadotte et al., 2009; Swenson et al., 2011; Robroek et al., 2017). Studies examining the relative importance of deterministic vs stochastic processes using randomized species and trait diversities are a relatively recent, and crucial fourth analytical advancement in characterizing DDRs (Swenson et al., 2011; Matthews et al., 2015; Si et al., 2016; Almeida-Gomes et al., 2019).

While there are a few studies that consider these concepts for taxonomic DDRs, there has been very little work that compares the relative contributions from turnover-vs-nestedness, environment-vs-geography, Quaternary-vs-contemporary climate and deterministic-vs-stochastic processes to both taxonomic and functional dissimilarity. Particularly lacking are studies that examine these multiple causative mechanisms within a comparative framework for different faunal taxa across a single elevational gradient. Majority of the studies on DDRs have investigated dissimilarities at large biogeographic scales, spanning multiple latitudes, using species presence / absence matrices (Poulin, 2003; Astorga et al., 2012; Wetzel et al., 2012; Fitzpatrick et al., 2013; but see Basset et al., 2015; Mori et al., 2015; González-Reyes et al., 2017; Tonkin et al., 2017). At large biogeographic scales, the ‘true’ environmental difference between communities is confounded by the added complexity of variation in historic climatic contingencies, which is seldom included in investigations (but see Fitzpatrick et al.., 2013).

In this study we compare the patterns and processes for the taxonomic and functional dissimilarity of ectothermic hawkmoths and endothermic birds across a single 2600m elevational transect in the eastern Himalaya of Arunachal Pradesh, India. The concurrent sampling of the two taxa, along the same elevational transect, is expected to reduce the number of confounding factors due to the identical parameters associated with climate, vegetation and history. We investigate the relative contribution of turnover and nestedness to both these facets of diversity. To assess the relative contribution of environment and geographic distance, we use the recently proposed Generalized Dissimilarity Modeling (GDM, Ferrier et al., 2007), and incorporate Quaternary climatic variables in the model as additional, independent predictors. We further ask the question whether the functional turnover between communities of birds and hawkmoths is higher or lower than expected, given the observed taxonomic turnover, i.e. randomness in trait distributions.

More specifically, we test the following hypotheses: (i) Due to the broad environmental gradient across a small spatial extent, we expect turnover to dominate both taxonomic and functional dissimilarities and, (ii) a strong positive correlation between the two facets of diversity for both organismal groups. Due to the difference in life histories for the two taxa (ectothermic hawkmoths versus endothermic birds) and their high vagility, we expect (iii) higher rate of turnovers for hawkmoths due to the positive association between body size and dispersal ability (Gaston & Blackburn, 1996; Soininen et al., 2018), and (iv) a higher relative contribution of temperature parameters to the observed beta diversity patterns of hawkmoths, as compared to birds.

## Materials and Methods

### Study site and field sampling

Light trapping for hawkmoths and transect counts for birds were carried out along the same transect in Eaglenest Wildlife Sanctuary (27.0–27.2°N, 92.3–92.6°E) in the state of Arunachal Pradesh, northeast India. The region, nested within the eastern Himalayan range, is one of the world’s 8 ‘*hottest hotspot*’ of biodiversity and endemism (Myers et al., 2000). Climate, vegetation and topography has been described elsewhere (Mungee & Athreya, 2019a).

Nocturnal phototropic Sphingidae were sampled at light screens at 13 elevations between 200 and 2800m. Methodology, rarefaction curves and species richness has been discussed elsewhere (Mungee & Athreya, 2019a). For birds, line transect surveys were conducted at a finer scale of 50m elevational resolutions. The counts were pooled within each 200m elevational band for comparisons with the elevational resolution of the hawkmoth data (see **Appendix S1**).

### Trait data sets

For hawkmoths, we used the morpho-functional traits of body mass, wing loading and wing aspect ratio (Mungee & Athreya, 2019a). Individual measurements for 3301 hawkmoths were obtained from field images after distortion-correction and size calibration (Mungee and Athreya, 2019b). For birds, we obtained species mean trait values for 227 (93%) out of the 245 birds in our sample from various sources (Dunning, 2008; Price et al., 2014; http://www.birdlife.org/). 6 quantitative and 3 categorical traits were used that have previously been linked to different avian functional strategies – body mass, wingspan, beak length, beak width, beak depth, tarsus length, primary substrate, foraging mode and diet (**Appendix S1**).

### Environmental data sets

Contemporary climate data was obtained from CHELSA climatologies (http://chelsa-climate.org/bioclim/), which is a recently assembled high resolution climatological data set that has been used in species distribution modeling with superior results (Karger et al., 2017). Rasters for mean annual temperature (MAT), maximum temperature of the warmest month (TMAX), minimum temperature of the coldest month (TMIN), annual precipitation (APPT) and precipitation seasonality (CVPPT) were downloaded at a 30m resolution and elevation specific mean values were obtained for the spatial coordinates of the sampling locations of hawkmoth light screens, which also correspond to the approximate mean elevations of each bird transect elevational band. The contemporary climatic data has been averaged from 1979 – 2013 (**Appendix S1**).

We assembled the same five bioclimatic variables (MAT, TMAX, TMIN, APPT and CVPPT) for the Last Glacial Maximum (∼ 22,000 years ago) using three commonly used Global Climate Models (GCMs) – Community Climate System Model 4 (CCSM4; Gent et al., 2011), Model for Interdisciplinary Research on Climate – Earth System Model (MIROC-ESM; Watanabe et al., 2011) and the Max Planck Institute – Earth System Model running in low resolution grid and paleo mode (MPI-ESM-P; Giorgetta et al., 2013). The Quaternary climatic stability was defined as the change in the contemporary and historic variable (Jansson, 2003) and was averaged across the three GCMs. We used an uncorrelated subset (r < 0.75) of contemporary climatic variables and Quaternary climatic stability to arrive at a final set of 5 environmental variables – TMAX-contemporary, APPT-contemporary, CVPPT-contemporary, Delta-MAT (change in MAT between LGM and present) and Delta-TMAX (change in TMAX between LGM and present). Only these five variables were used for all subsequent analyses.

### Statistical analyses

#### Taxonomic and functional dissimilarity

Taxonomic dissimilarity was calculated using the abundance-based Bray-Curtis dissimilarity index (Baselga, 2013). For functional dissimilarities, the species-by-trait matrix was converted into a distance matrix using Podani’s extension for ordinal traits (Podani 1999). The distance matrix was used for cluster analysis (UPGMA method) to create a dendrogram (Petchey & Gaston 2002), which was subsequently converted to a functional tree. The abundance weighted SØrensen dissimilarity index was used to generate pair-wise dissimilarities across all sites. To quantify the relative importance of species turnover and nestedness to the overall dissimilarity, we used the procedures described by Baselga (2010, 2013) for abundance-based Bray-Curtis dissimilarity and extended for SØrensen (functional) dissimilarity by Leprieur et al. (2012). We also checked for correspondence between the taxonomic dissimilarities generated using Bray-Curtis index and SØrensen index using Mantel test.

#### Generalized Dissimilarity Modeling

To evaluate the relative contributions of environmental and geographic distances, we used Generalized Dissimilarity Modeling (GDM; Ferrier et al., 2007). GDM is a non-linear extension to matrix regression which can (i) account for the curvilinear relationships between community dissimilarity and environmental/geographic distances, (ii) assess the independent roles of multiple predictors, (iii) account for the variation in the strength of the relationship (between dissimilarity and individual predictor) along the gradient, and (iv) be used for model-deviance-partitioning (Borcard et al., 1992) to calculate the joint and independent contribution of geographic and environmental distances (Fitzpatrick et al., 2013).

For each predictor, GDM performs a transformation using a set of I-spline functions. The I-splines are essentially short stretches of polynomial functions, ‘stitched’ together with a high degree of smoothness. The coefficients for each I-spline are determined using maximum likelihood estimation and the model standardizes the different predictors to allow for a direct comparison. We used the default of three I-spline basis functions per predictor (Fitzpatrick et al., 2013). Each function gives two important pieces of information regarding the relationship between the predictor and the dissimilarity – (i) the maximum height of the I-spline function, i.e. the sum of the three coefficients, is an estimate of the proportion of turnover explained by that predictor, and (ii) the difference in the height of the function between any two points along the gradient, describes the variation in the relationship between the predictor and the dissimilarity. We fit the GDMs using both – the taxonomic and the functional dissimilarity matrix for hawkmoths and birds. We refer to these as taxonomic-GDM and functional-GDM below.

For both dissimilarities, taxonomic and functional, three separate GDMs were fitted to assess the relative contribution of environment and space - (i) full model – with both sets of predictors (environmental and and geographic), (ii) only environmental predictors (both contemporary and historic), and (iii) only geographic distances. The full model deviance was partitioned into these three independent components using the variation partitioning method of Borcard et al. (1992). For significance testing of variables and model selection, we performed Monte Carlo sampling (999 permutations) and step-wise backward elimination. Relative importance of each predictor was obtained from the scaled values of the maximum height of the corresponding I-spline functions.

#### Null Modeling Analysis

A null distribution of functional beta diversity values was generated for each trait, and for overall functional dissimilarity using all traits, by randomizing (999 times) the names of the species across the tips of the trait dendrograms. Therefore, the randomization procedure maintains the species richness, relative abundance distributions and consequently taxonomic beta diversity at each elevational community (Swenson, 2011). A standardized effect size (SES) was calculated for functional beta diversity using the mean and standard deviation of the null distribution as follows: 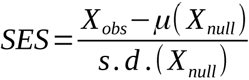, where *X*_*obs*_ is the observed dissimilarity value between two communities, *μ(X*_*null*_) the mean of the null distribution and *s.d.(X*_*null*_) the standard deviation of the null distribution. Values greater than 1.96 indicate a higher than expected functional dissimilarity between the communities and *vice versa*.

We additionally calculate functional redundancy (FRed) metric as a measure of resilience of hawkmoth and bird communities to environmental change across the elevational gradient. FRed was defined as the difference between Simpson’s species diversity and functional diversity, and ranges from 0 to 1, indicating complete divergence or convergence between the two facets, respectively (Robroek et al., 2017). Observed FRed for the hawkmoth and bird communities was compared with those obtained from 999 randomly assembled matrices, using SES values as previously. We also compared Simpson’s and SØrenson’s functional dissimilarities using ordinary least squares regression.

Finally, to compare the results from GDM with an analogous non-parametric linear regression approach, we used partial mantel tests and its more commonly used extension – distance based multiple matrix regressions with randomizations (MMRR; Wang 2013). Correlation between taxonomic dissimilarities (or functional dissimilarities) and environmental distances was obtained while accounting for geographic distances, and *vice versa*. The significance of the statistics was assessed with 999 permutations.

All analyses were performed in R 3.4.4 on a Ubuntu (linux-gnu) 18.04.1 platform (R Core Development Team 2013). Individual functions and packages used for various analyses have been provided as **Appendix S2**.

## Results

### Taxonomic and functional dissimilarity

We recorded a total of 4,731 hawkmoth individuals spanning 80 morphospecies, 30 genera and 3 subfamilies. We reliably measured morpho-functional traits of body mass, wing loading and wing aspect ratio for 3,301 individuals and arrived a species mean trait values for all species (Mungee & Athreya 2019a, 2019b). 15,387 individual birds, spanning 235 species, 150 genera and 50 families were recorded and species mean traits were obtained from various published sources for a subset of 227 (93%) of the species (Dunning, 2008; Price et al., 2014; http://www.birdlife.org/) (**Appendix S1**).

Functional and taxonomic dissimilarities were strongly correlated (*hawkmoths* – *Mantel’s r = 0.84, p* < *0.005; birds – Mantel’s r = 0.93, p* < *0.005;* **Figure 1**). The relationship was linear for hawkmoths (β_func_ *- 0.30/β*_*taxo*_ + *0.05, r*^*2*^ *= 0.70, p* < *0.005*) but the quadratic relationship was a superior fit for birds; ΔAIC > 20 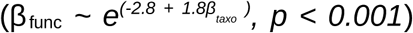. Bray-Curtis and SØrensen taxonomic dissimilarities were highly correlated (hawkmoths – Mantel’s r = 0.92, p < 0.001; birds – Mantel’s r = 0.96, p < 0.001; **Appendix S2**). The taxonomic beta diversity was dominated by species turnover (92 % for hawkmoths and 97 % for birds), while nestedness component had a considerable contribution to the functional beta diversity (31 % for hawkmoths and 24 % for birds) **(Figure 1)**. Overall, bird communities exhibited a greater slope for both taxonomic and functional DDRs than hawkmoth communities (taxonomic dissimilarity Fisher’s z = 8.68, p.value < 0.001; functional dissimilarity Fisher’s z = 6.03, p.value < 0.001; **Appendix S2**). The taxonomic dissimilarities for birds using the subset of 227 species (for which functional dissimilarities were calculated) exhibited similar slopes (**Appendix S2**).

**Figure 1.**
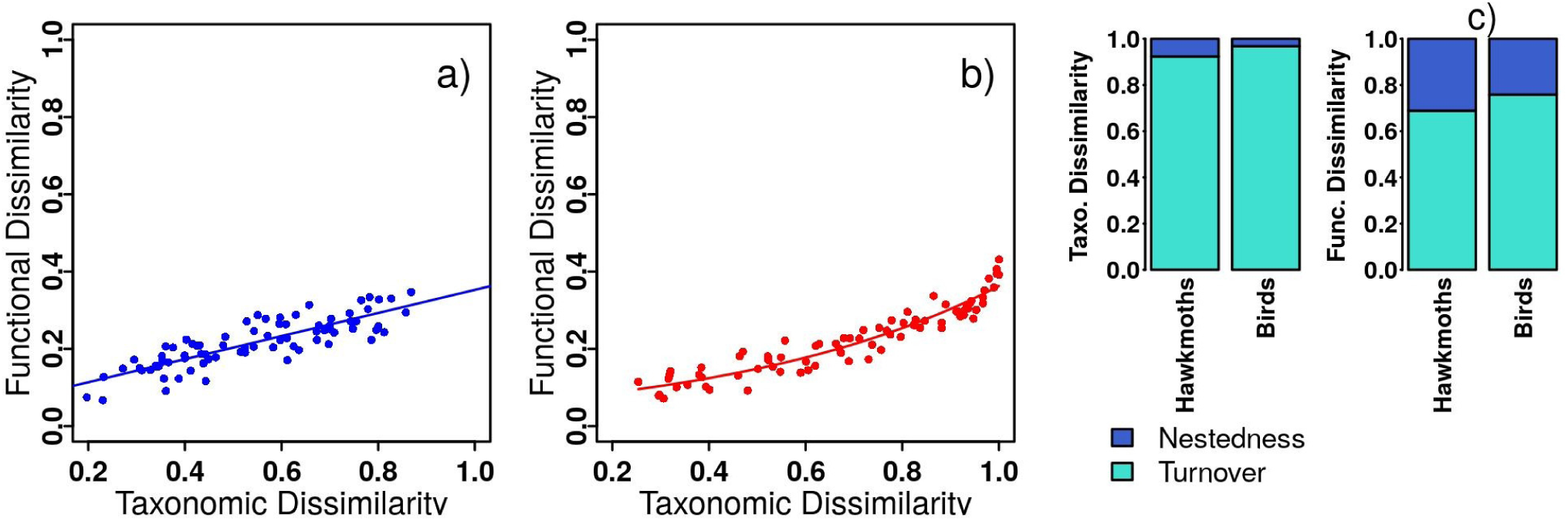
Relationship between functional and taxonomic dissimilarities for communities of a) Hawkmoths and b) Birds; c) The proportion of total dissimilarity attributable to the turnover and nestedness components for taxonomic (left) and functional (right) beta diversity for hawkmoths and birds.

### Generalized Dissimilarity Modeling

For taxonomic-GDM, 90% deviance could be explained by using the full model (environmental and geographic distances both) for birds, whereas deviance explained for hawkmoths was slightly lower (80%). Joint and independent effects of geographic distance and environment varied between hawkmoths and birds with over 50% contribution attributed to purely environment for hawkmoths as compare to 31% for birds, whereas the joint contribution of environment and geographic distance was much higher for birds (47%) than hawkmoths (10%). Geographic distance alone explained very little deviance for hawkmoths and birds (19% and 11%, respectively). The relative importance of individual predictors differed strongly for taxonomic dissimilarities of the two organismal groups. Hawkmoth communities were more strongly correlated with contemporary climate, especially MAT (52%) and APPT (24%), whereas bird communities exhibited strongest correlations with Quaternary climatic changes, especially Delta-MAT (60%) (**Figures 2 & 3; Table 1**).

**Table 1.** The proportion of total explained deviance attributable purely to space, purely to environment, jointly to both variables (shared), and not explained by the fitted GDM for the taxonomic and functional dissimilarities of hawkmoths and birds across the elevational gradient. The independent, relative contributions from the individual predictors, while keeping all others constant, are also shown.

|  |  | Unexplained | Spatial + Env. | Spatial | Environmental |
| --- | --- | --- | --- | --- | --- |
| Taxonomic | Hawkmoths | 0.20 | 0.10 | 0.20 | 0.51 |
|  | Birds | 0.11 | 0.47 | 0.11 | 0.31 |
| Functional | Hawkmoths | 0.34 | 0.34 | 0.00 | 0.31 |
|  | Birds | 0.11 | 0.50 | 0.12 | 0.28 |

|  |  | Unexplained | Spatial | MAT -<br>contemporary | APPT -<br>contemporary | CVPPT -<br>contemporary | Delta -<br>MAT | Delta -<br>CVPPT |
| --- | --- | --- | --- | --- | --- | --- | --- | --- |
| Taxonomic | Hawkmoths | 0.20 | 0.02 | 0.52 | 0.24 | 0.02 | 0.00 | 0.00 |
|  | Birds | 0.11 | 0.18 | 0.11 | 0.00 | 0.00 | 0.60 | 0.00 |
| Functional | Hawkmoths | 0.34 | 0.01 | 0.33 | 0.23 | 0.08 | 0.00 | 0.00 |
|  | Birds | 0.11 | 0.16 | 0.02 | 0.22 | 0.00 | 0.41 | 0.07 |

**Figure 2.**
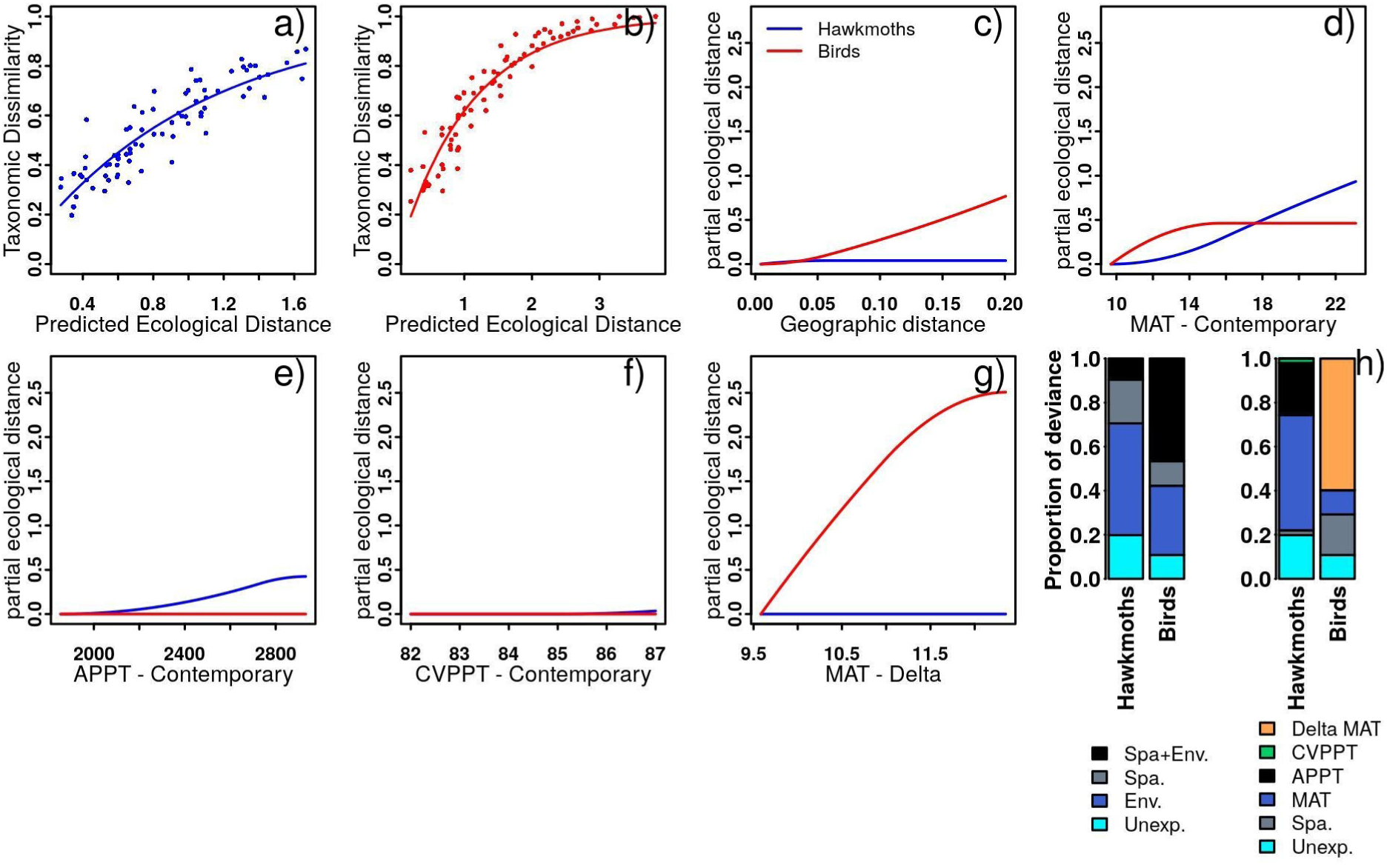
Relationship between observed functional dissimilarity of each elevational-site pair for a). hawkmoths, and b). birds, with the fitted predictor from the GDM (predicted ecological distance between elevational-site pairs); Partial regression fits (I-splines) for the different predictors significantly associated with either hawkmoth (blue) or bird (red) functional dissimilarities are shown in plots c) through g). The maximum height and shape of each function provides an indication of the independent contribution of the predictor and variation in it’s strength along the environmental gradient. Relative importance of each predictor, obtained from the scaled values of the maximum height of the corresponding I-spline functions is shown in h).

**Figure 3.**
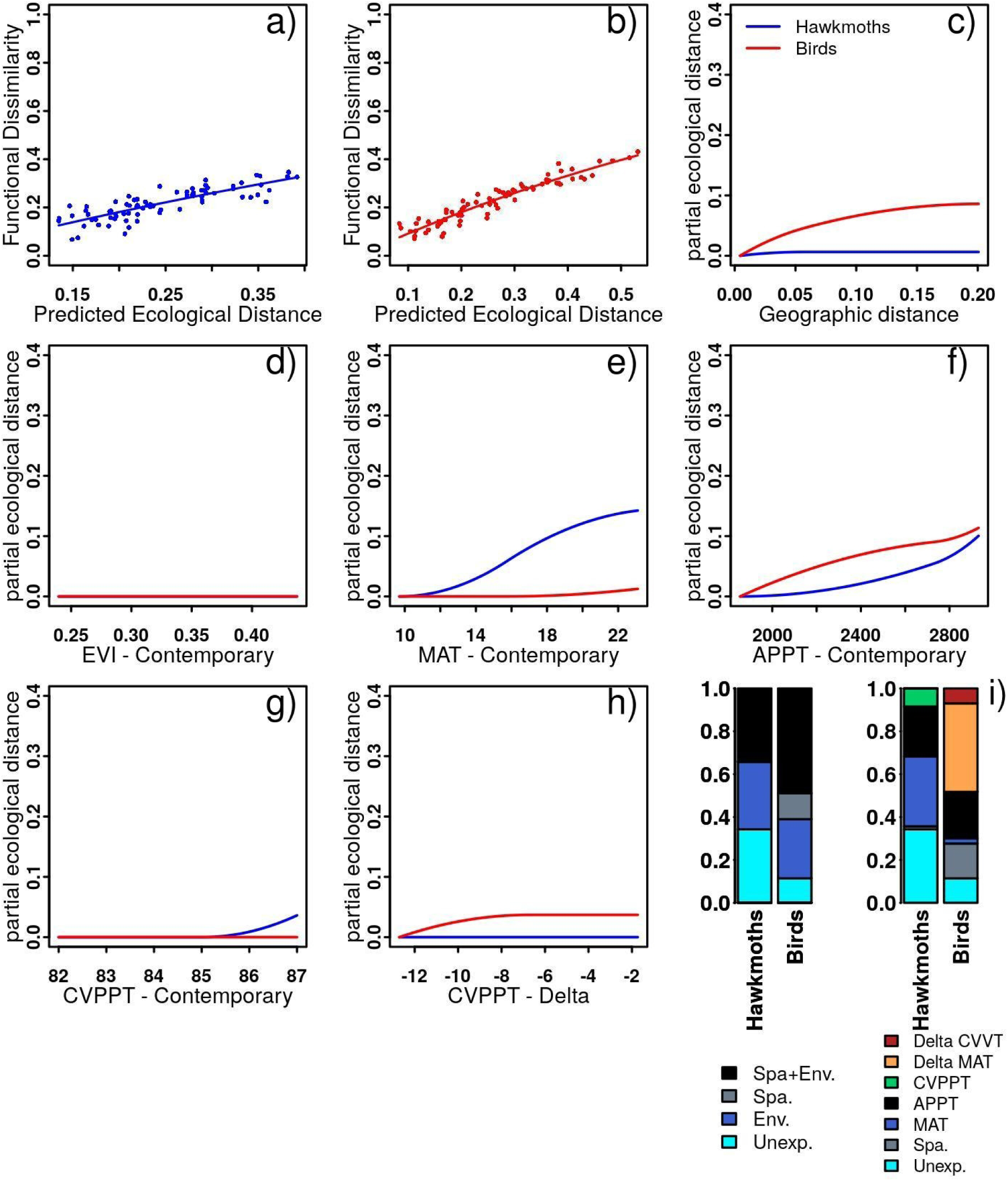
Relationship between observed functional dissimilarity of each elevational-site pair for a). hawkmoths, and b). birds, with the fitted predictor from the GDM (predicted ecological distance between elevational-site pairs); Partial regression fits (I-splines) for the different predictors significantly associated with either hawkmoth (blue) or bird (red) functional dissimilarities are shown in plots c) through h). The maximum height and shape of each function provides an indication of the independent contribution of the predictor and variation in it’s strength along the environmental gradient. Relative importance of each predictor, obtained from the scaled values of the maximum height of the corresponding I-spline functions is shown in i).

Similar to taxonomic-GDM, the functional-GDM for hawkmoth communities was more strongly correlated with contemporary climate, especially MAT (33%) and APPT (23%), whereas bird communities exhibited strong correlations with Quaternary climate, especially Delta-MAT (41%). Only 64% deviance could be explained by using the full model (environmental and geographic distances both) for hawkmoths, whereas deviance explained for birds was close to 90% for functional dissimilarities. Variance partitioning showed that the relative contribution by the environment alone was similar for hawkmoths (31%) and birds (28%), whereas the contribution of spatially limited dispersal was higher for birds (12%) than hawkmoths (0%) (**Figures 2 & 3; Table 1**).

### Null Modeling Analysis

SES values for the taxonomic beta diversity were significantly greater than null for both hawkmoths and birds (except one high elevation community for hawkmoths; **Figure 4)**. Additionally, the SES values exhibited a significant negative slope with elevation for hawkmoth taxonomic dissimilarity. Interestingly, the overall functional beta dissimilarity across all traits was not significantly different from null for majority of the communities of hawkmoths and birds. The patterns were very similar using individual traits for hawkmoths and birds, and are presented in **Appendix S2**. The observed functional redundancy values were high for both organismal groups (Hawkmoths - FRed_μ_ = 0.55, FRed_σ_ = 0.02; Birds – FRed_μ_ = 0.51, FRed_σ_ = 0.02 **Appendix S2**). The values were not significantly different from null at most elevations, except the highest elevations for birds (**Figure 4**).

**Figure 4.**
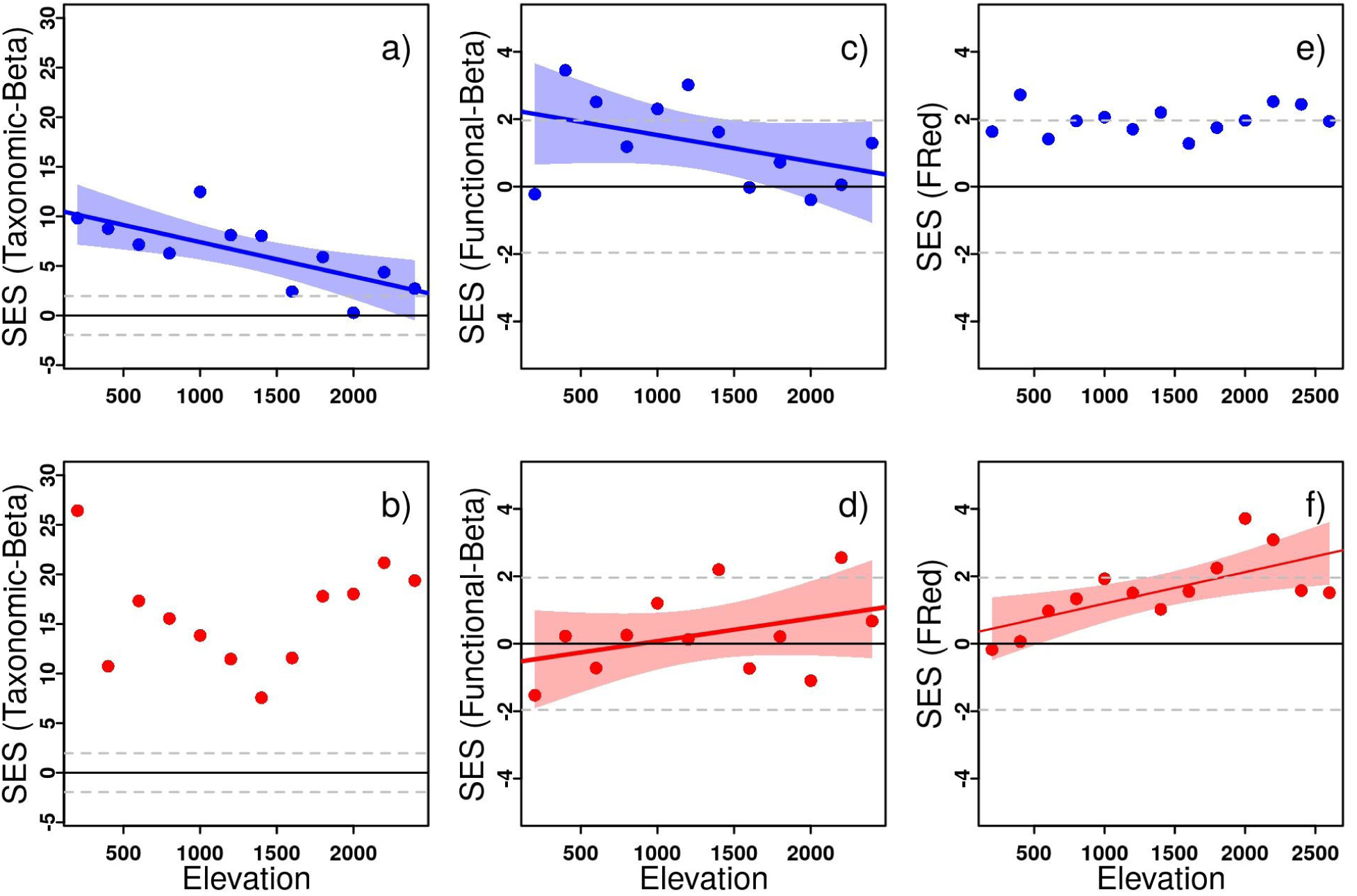
SES values for the deviation of observed metrics of taxonomic beta-diversity (a & b), functional beta-diversity (c & d) and Functional redundancy (e & f) along the elevational gradient, presented for hawkmoths (blue; top row) and birds (red; bottom row). SES values greater than 1.96, or less than −1.96 (the dashed grey lines) indicate that the observed values of the respective metrics are higher or lower than the values obtained under randomized assembly (see text for details).

Partial Mantel tests showed similar results for birds, with a higher contribution of geographic distances to taxonomic and functional dissimilarities (Mantel’s R_taxonomic_ = 0.61, p.value < 0.005; Mantel’s R_functional_ = 0.66, p.value < 0.005) as compared to the independent contribution from environment alone (Mantel’s R_taxonomic_ = 0.20, p.value = 0.06; Mantel’s R_functional_ = 0.14, p.value = 0.17). However, for hawkmoths results were different from GDM, indicating little contribution of environment (Mantel’s R_taxonomic_ = 0.11, p.value = 0.18; **Figure** Mantel’s R_functional_ = 0.12, p.value = 0.18) as compared to geographic distance (Mantel’s R_taxonomic_ = 0.48, p.value < 0.005; Mantel’s R_functional_ = 0.29, p.value < 0.05). Results were similar with MMRR; environmental distance did not show significant correlation with either of the diversity facets for the two taxa, whereas geographical distances were strongly correlated with both facets of dissimilarity and with both taxa. Similar to GDM, the coefficients for the overall MMRR model (including geographic distance and environmental distance matrices) were higher for birds (R^2^_taxonomic_ = 0.60, p.value < 0.05; R^2^_functional_ = 0.63, p.value < 0.05) than hawkmoths (R^2^_taxonomic_ = 0.41, p.value < 0.05; R^2^_functional_ = 0.21, p.value < 0.05) (**Appendix S2**).

## Discussion

We compared the patterns and processes for taxonomic and functional dissimilarities of hawkmoths and birds across a broad elevational gradient in the east Himalayan global biodiversity hotspot. The two facets of diversity exhibit strong correlation, however, despite high species turnover, the functional composition of the communities exhibited high redundancy and higher nestedness. The high randomness in the distribution of traits across communities may be indicative of community resilience to changing environment with the overall community-level functional roles remaining constant. Overall, the two facets of diversity had very similar relative contributions from the abiotic predictors for a given organismal group, but there was significant variation across taxa.

### Taxonomic and functional dissimilarities

Our results show that the further apart two sampling sites are, along an elevational gradient, the more dissimilar they are in terms of both species and functional composition, however the contributions of turnover-versus-nestedness varied considerably across the two facets of diversity. There was higher functional nestedness across both taxa, as compared to taxonomic nestedness, indicating that the functionality of local communities are increasingly nested subsets of the total suite of available functionalities in the regional pool, in spite of a high species turnover. Contrary to our predictions, we observed a significantly higher rate of turnover for birds as compared to hawkmoths, which gives valuable taxon-specific signatures (slope of the DDR; **Appendix S3**).

Hawkmoths and birds are both active dispersers, and their range size may be expected to increase with body size (Gaston & Blackburn, 1996). Contrary to this expectation, we observed a significantly higher rate of turnover for birds. This result supports the most recent meta-analysis on the subject – body size & beta-diversity relationships do not follow a universal trend and may be context dependent (Soininen et al., 2018). Many tropical bird species are highly specialized, exhibit high endemism, narrow niche widths and extreme dispersal limitation (Moore et al., 2008 and references therein). Unlike the tropical bird species, hawkmoth species are found throughout the Indo-Australian archipelago and have demonstrated a very broad resource utilization spectrum (Beck et al.., 2006; Beck et al., 2007).

As with any functional diversity related analysis, a comment of the implications of the traits used is warranted. The hawkmoth morphological traits used here exhibit a strong response to the environmental gradient in the study region (Mungee & Athreya 2019a) and have previously been implicated in resource requirements, thermoregulation and dispersal ability (Heinrich 1996, Hassal 2015, Vágási et al. 2016). However larval diet breadth is an important strategy that has shown to correlate strongly with the distribution of Sphingidae across the Indo-Malayan Archipelago (Beck et al., 2006; Beck & Kitching, 2007). The traits used for bird assemblages on the other hand, encompass broader categories of functionality across resource use, thermoregulation, dispersal ability, feeding guild or the impacts of species on other trophic levels (Petchey et al., 2007; Flynn et al., 2009; Ding et al., 2013; Price et al., 2014; Pigot et al., 2016), and thus may better encapsulate the functional dissimilarities across large distances where communities exhibit largely non-overlapping taxonomic compositions (Bray-Curtis dissimilarities > 0.97). Due to a lack of information on the hawkmoth host plants and their distribution in the study region, currently we do not have the means to account for their resource niches.

There are only a few analyses of changes in functional composition of animal communities along tropical altitudinal gradients, and there are still large gaps in knowledge regarding the role of functional beta diversity in maintaining ecosystem resilience of tropical assemblages (Villéger et al., 2013; Dehling et al., 2014; Nunes et al., 2016). Our findings indicate high redundancy in traits, which may be an important mechanism for tropical ecosystems to retain a fundamental, base-line functionality despite the high rate of species turnover (Mori et al., 2015).

### Generalized Dissimilarity Modeling

The ecological literature is replete with several predictions for the variation in distance dissimilarity relationships across regions and across taxa (Nekola & White, 1999; Palmer, 2005; Soininen et al., 2007; Soininen et al., 2018). On the contrary, there are very few general hypotheses for determining the relative role of geographic and environmental distances and most meta-analyses indicate idiosyncratic, taxon-specific contributions of these two non-mutually exclusive predictors (Fitzpatrick et al., 2013; Glassman et al., 2017; KÖnig et al., 2017). Using Generalized Dissimilarity Modeling, we demonstrated that rates of compositional dissimilarity vary substantially as a function of the predictor considered and with position along gradients, thus enabling the identification of regions of high vulnerability associated with different predictors in isolation and in quantifying the sensitivity of different ecological communities to future perturbations. Beyond indicating greater overall turnover for the birds of tropical Himalaya than the hawkmoths, the functions from GDM suggested that the historic environmental gradients, especially mean temperature, and spatially limited dispersal, most strongly associated with the beta diversity of birds, whereas the hawkmoths distribution was shaped by contemporary climate (mean temperature and annual precipitation). Thus, while MAT was the single best predictor for beta diversity patterns of both hawkmoths and birds, the relative importance of contemporary and historic temperatures was widely disparate.

A primary result from the GDM analysis was the similarity in the relative contributions of different predictors across the two facets of diversity for the same organismal group. This is interesting and indicates that while there is a large difference in the rate of response of taxonomic and functional turnover of a taxa, to the same environmental gradient, their relationship with individual predictors is similar. Apart from the higher unexplained variance in functional-GDM of hawkmoths, contemporary mean annual temperature had the highest relative contribution to both facets of dissimilarity, followed by contemporary precipitation. Historic climate did not contribute at all to the observed taxonomic and functional dissimilarities of hawkmoths of eastern Himalaya. Delta-MAT, i.e. the change in the mean annual temperature between the Last Glacial Maximum and present, was the most important correlate for both – taxonomic and functional dissimilarity of birds. The joint and independent contributions of environmental and geographic distance was remarkably similar across the two facets of dissimilarities for birds (Joint > Env. > Spatial), whereas it was slightly more variable for hawkmoths. The discrepancy between the relative contributions of individual predictors was however magnified when comparing across taxa, even for the same facet of dissimilarity.

Overall, our results suggest that variation in the relative role of environmental and geographic filters in determining beta-diversity patterns is persistent across organismal groups even along the same, identical elevational transect. While the drivers of beta-diversity exhibit idiosyncrasy and taxon-specificity; for a given taxa, they are consistent across the two facets of dissimilarity. The consistency of this pattern, across two disparate organismal groups, is suggestive of a key mechanism in which tropical communities may retain functionality of ecosystems in a changing environment. More importantly, regardless of the principal predictor, the net result was that of high taxonomic turnover, which is de-coupled from functional turnover for two contrasting taxa. The large redundancy in trait values, despite high species turnover, indicates functional resilience of these tropical communities. Such comparative studies on the relationship of different environmental predictors, across multiple facets of diversity will help improve our understanding of the processes generating beta-diversity in the species rich tropical systems.

## Acknowledgments

We thank the Forest Department of Arunachal Pradesh for their assistance and research permits (CWL/G/13(95)/2011-12/Pt.II/660-62 during 2011-2015). We are especially grateful to Mr. Millo Tassar, the DFO (Shergaon Forest Division), for his tremendous help and encouragement. We thank all the members of the Singchung Bugun community for their support. The project was partly funded by the Department of Science and Technology, Government of India (Grant No. SR/SO/AS66/2011) and by the Nadathur Trust, Bengaluru. MM would like to acknowledge the comments of Amod Zambre, which benefited this manuscript. The authors would also like to thank Dr. L. S. Shashidhara (IISER, Pune) for his fostering confidence.

## Data availability Statement

The data has been provided as Online Supporting Information.

## Biosketch

**Mansi Mungee** is a Postdoctoral fellow at the Wildlife Institute of India, Dehradun. She has recently submitted (June, 2019) her doctoral thesis at the Indian Institute of Science Education and Research (IISER-Pune). Her main interests are community ecology, functional traits and tropical biodiversity. She is particularly interested in the functional trait ecology of eastern Himalayan lepidoptera.

**Ramana Athreya** is an Associate Professor (Physics & Biology) at IISER-Pune, India. With a formal training in astronomy, he has also been working for the conservation of forests in Arunachal Pradesh for the last 15 years, His principal interests in biology are diversity research (diversity patterns and speciation processes) and wildlife conservation paradigms for Arunachal Pradesh.

