## Appendix for "Functional randomness despite high taxonomic turnover across an elevational gradient in a global biodiversity hotspot: A case study of hawkmoths and birds"

**Appendix S1**

**Online Supporting Information**

Functional randomness despite high taxonomic turnover across a broad elevational gradient in a global biodiversity hotspot: A case study of hawkmoths and birds

### Field sampling and data

Details for hawkmoth sampling and trait measurements can be obtained from Mungee & Athreya 2019a & 2019b. All bird sampling was conducted by a single person (Rohan Pandit) to minimize identification bias. Forty-nine elevations between 200 and 2800 m, in intervals of 50 m were surveyed long the Eaglenest road. Birds were recorded along a 200m transect during a steady walk over 5 minutes. The same transect was sampled in two sets, with a steady walk in both directions – to and fro. The two counts were recorded separately. Birds recorded, both visually and aurally, within a perpendicular distance of approximately 20m from the road were dictated into a voice recorder. These audio clips were later transcribed into a spreadsheet and checked for errors by two different transcribers. The same transects were sampled on 12 different days (12 sets per elevation; May and June, 2012-14). The sampling was done during 6.00-12.00 hr. A maximum of 12 elevations were covered during a day, though rain often reduced the number. We avoided any systematic bird activity difference across the large 6-hr window, by subdividing it into three 2-hr slots – Early morning (E), Mid morning (M) and Late morning (L). The 12 transects at each elevation were equally distributed across these 3 slots. Security issues precluded us from sampling elevations below 500m inside the sanctuary, and the 200m elevation transects were conducted in the neighbouring Pakke Tiger Reserve, about 20 km away from the 500m transect. It should be however noted that we could not avoid the major barrier – the gorge of the Kameng river – between these two lowermost elevations.

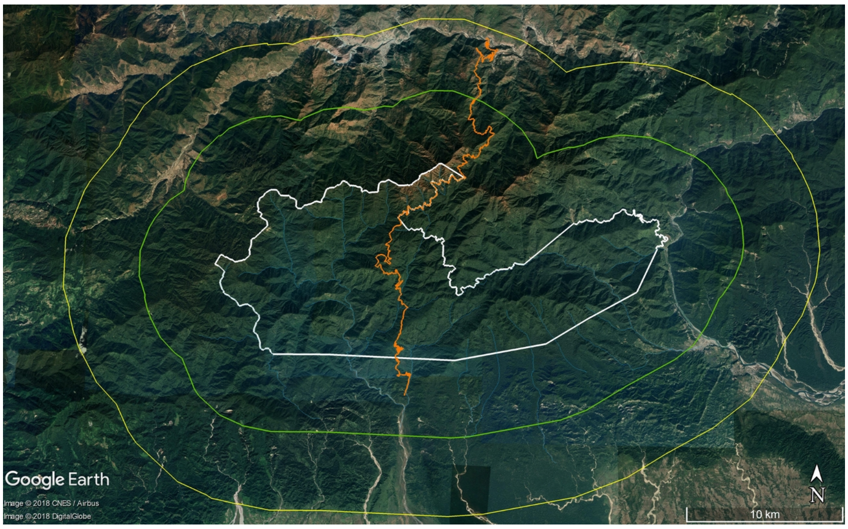


**Figure S1.** Study site (Eaglenest Wildlife Sanctuary; boundary shown in white above with contours for 5 & 10km buffer marked in green and yellow respectively) was located in northeast India. The bird transects and the hawkmoth sampling was conducted along the Eaglenest road (shown in organe). The lowest elevation sampling (i.e. 200m) was situated outside the sanctuary and is markes in red for birds and blue for hawkmoths. (Also see Figure1; Mungee & Athreya 2019a).

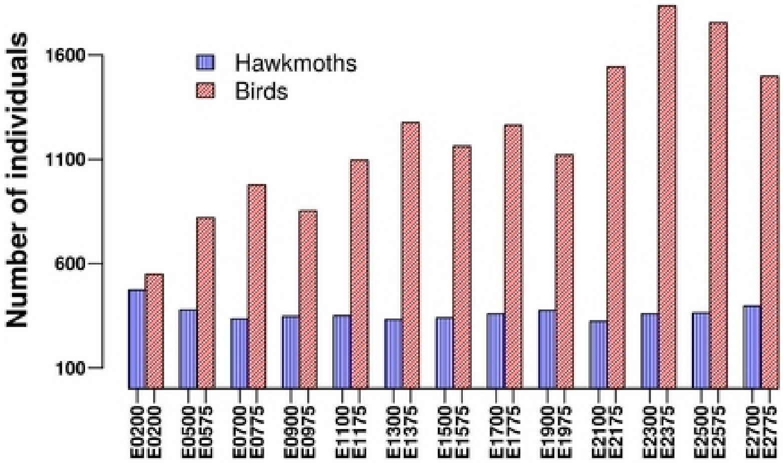


**Figure S2**. The number of individuals recorded at various elevations for hawkmoths (blue) and birds (red). The bird counts were pooled across four 50m elevational bins to obtain a 200m elevational resolution.

**Table S1.** Summary statistics for the hawkmoth and the bird data. N = number of individuals recorded, Sobs = Number of species observed, Srare = Rarefied species richness; Alpha = Species diversity calculated using Fisher's alpha diversity index. The row names correspond to the elevation code (e.g. E0200 = 200m sampling elevation).

| | **N** | | **$S_{obs}$** | | **$S_{rare}$** | | **Alpha** | |
|---|---|---|---|---|---|---|---|---|
| | **Hawkmohts** | **Birds** | **Hawkmohts** | **Birds** | **Hawkmohts** | **Birds** | **Hawkmohts** | **Birds** |
| **E0200** | 473 | 549 | 43 | 53 | 39 | 53 | 11.5 | 15 |
| **E0500** | 378 | 820 | 39 | 81 | 37 | 74 | 10.9 | 22 |
| **E0700** | 335 | 977 | 33 | 84 | 33 | 76 | 09.1 | 22 |
| **E0900** | 347 | 852 | 45 | 77 | 44 | 70 | 13.8 | 21 |
| **E1100** | 350 | 1095 | 48 | 88 | 46 | 76 | 15.1 | 23 |
| **E1300** | 332 | 1276 | 31 | 82 | 31 | 73 | 08.4 | 20 |
| **E1500** | 340 | 1162 | 40 | 87 | 39 | 78 | 11.8 | 22 |
| **E1700** | 359 | 1264 | 36 | 85 | 35 | 74 | 10.0 | 21 |
| **E1900** | 376 | 1120 | 40 | 77 | 38 | 68 | 11.3 | 19 |
| **E2100** | 323 | 1543 | 29 | 73 | 29 | 63 | 07.7 | 16 |
| **E2300** | 359 | 1835 | 27 | 81 | 26 | 66 | 06.8 | 17 |
| **E2500** | 363 | 1755 | 23 | 77 | 22 | 62 | 05.5 | 17 |
| **E2700** | 396 | 1498 | 32 | 69 | 30 | 56 | 08.2 | 15 |
| **Total** | **4731** | **15746** | **80** | **245** | **----** | **----** | **----** | **---** |

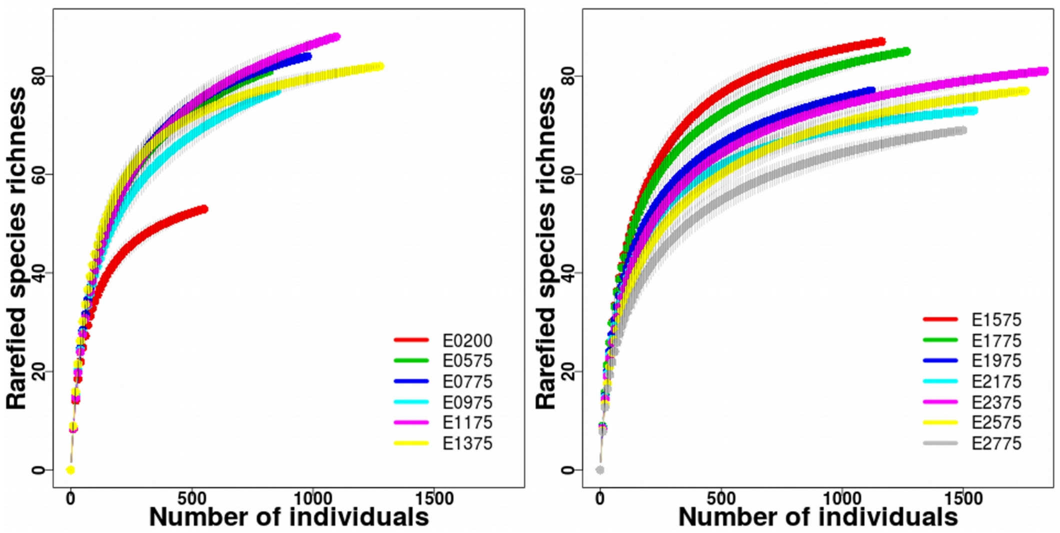


**Figure S3**. Rarefaction curves for the individual elevational community of birds.

**Table S2.** Bird species list and trait data set used for all the analyses. All measurements are in mm. *Blength = Beak length; Bwidth = Beak width; Bdepth = Beak depth; Wing = Wing length, Tarsus = length from the base of tarsus to last scute; PySub = Primary substrate (M = mid-canopy, T = canopy, B = bush, G = ground); Forag = Foraging mode (L = leaf, A= air, M = moss, G = ground, F = fruiting bodies, Fl = flower, B = bark, Ba = Bamboo); Diet = primary diet of the species during breeding season (I = insectivore, F = Frugivore, S = seeds, N = nectarivorous, O = Omnivore/Ophiophagus, V = Vertebrates); Mass = Body mass (in grams) (See Price et al., 2014 for further details on bird trait measurements).*

| | Blength | Bwidth | Bdepth | Wing | Tarsus | Pysub strate | ForagingMode | Diet | Mass |
|---|---|---|---|---|---|---|---|---|---|
| *Aegithalidae Aegithalos concinnus* | 4.9 | 2.53 | 3.25 | 52.5 | 17.2 | M | L | I | 6.1 |
| *Aegithinidae Aegithina tiphia* | 10.63 | 3.69 | 4.02 | 66.4 | 19.15 | T | L | O | 14.4 |
| *Alcedinidae Halcyon coromanda* | 42.04 | 14.93 | 14.47 | 106 | 15.81 | M | R | V | 77.5 |
| *Bucerotidae Aceros nipalensis* | 187 | 50.78 | 76.3 | 455 | 70.51 | T | F | F | 2385 |
| *Bucerotidae Anthracoceros albirostris* | 126.94 | 37.61 | 79.83 | 300.5 | 59.72 | T | F | F | 786.8 |
| *Bucerotidae Buceros bicornis* | 268 | 55.96 | 120 | 532.5 | 87.89 | T | F | F | 2798.5 |
| *Bucerotidae Rhyticeros undulatus* | 183 | 51.79 | 81.07 | 480 | 67.98 | T | F | F | 2232.5 |
| *Campephagidae Lalage melaschistos* | 12.76 | 6.23 | 6.07 | 118.5 | 21.81 | T | L | I | 43 |
| *Campephagidae Pericrocotus brevirostris* | 7.35 | 4.05 | 3.93 | 86.8 | 15.28 | T | L | I | 16.5 |
| *Campephagidae Pericrocotus ethologus* | 8.16 | 4.12 | 3.97 | 92.3 | 17.2 | T | L | I | 19 |
| *Campephagidae Pericrocotus flammeus* | 12.07 | 5.92 | 6.32 | 107 | 19.48 | T | L | I | 21.8 |
| *Campephagidae Pericrocotus solaris* | 8.07 | 4.03 | 4.22 | 84 | 16.36 | T | L | I | 14.5 |
| *Certhiidae Certhia discolor* | 10.4 | 2.45 | 2.45 | 70.5 | 17.55 | M | B | I | 10.4 |
| *Chloropseidae Chloropsis aurifrons* | 16.01 | 5.01 | 5.21 | 96.4 | 19.44 | T | L | O | 32.5 |
| *Chloropseidae Chloropsis hardwickii* | 15.69 | 5.01 | 5.71 | 95.8 | 19.32 | T | L | O | 32.8 |
| *Cisticolidae Orthotomus sutorius* | 9.41 | 2.73 | 2.54 | 46 | 19.96 | B | L | I | 7.5 |
| *Columbidae Chalcophaps indica* | 11.77 | 3.61 | 4.42 | 150.8 | 26.27 | G | G | S | 121 |
| *Columbidae Columba pulchricollis* | 12.98 | 5.6 | 5.68 | 203.1 | 25.37 | T | F | F | 330 |
| *Columbidae Ducula aenea* | 17.56 | 7.78 | 6.62 | 232 | 31.34 | T | F | F | 545 |

| | | | | | | | | | |
|---|---|---|---|---|---|---|---|---|---|
| *Columbidae Ducula badia* | 16.97 | 9.02 | 8.43 | 240.3 | 28.94 | T | F | F | 486 |
| *Columbidae Macropygia unchall* | 11.08 | 4.26 | 4.97 | 197.6 | 26.31 | T | G | F | 168 |
| *Columbidae Treron apicauda* | 11.57 | 5.97 | 6.92 | 161 | 22.77 | T | F | F | 220 |
| *Columbidae Treron curvirostra* | 11.2 | 5.53 | 7.95 | 140.5 | 21.33 | T | F | F | 134.5 |
| *Columbidae Treron sphenurus* | 11.58 | 6 | 6.28 | 178 | 23.33 | T | F | F | 210 |
| *Corvidae Cissa chinensis* | 25.6 | 11.34 | 13.87 | 148.8 | 46.29 | M | L | I | 126.8 |
| *Corvidae Dendrocitta formosae* | 22.75 | 11.79 | 14.31 | 138.8 | 30.12 | T | F | O | 94 |
| *Corvidae Dendrocitta frontalis* | 21.24 | 11.48 | 14.65 | 133.8 | 29.84 | T | F | O | 97 |
| *Corvidae Nucifraga caryocatactes* | 35.96 | 14.84 | 15.62 | 218.1 | 43.08 | T | F | S | 197.1 |
| *Corvidae Urocissa flavirostris* | 24.37 | 12.44 | 13.34 | 190 | 49.01 | T | L | I | 150.5 |
| *Cuculidae Cacomantis merulinus* | 13.02 | 5.45 | 4.99 | 115.5 | 20.95 | T | L | I | 25.3 |
| *Cuculidae Cacomantis sonneratii* | 9.89 | 6.76 | 5.64 | 123.9 | 16.07 | T | L | I | 33.8 |
| *Cuculidae Chrysococcyx maculatus* | 12.41 | 5.29 | 5.06 | 107.5 | 19.59 | T | L | I | 24 |
| *Cuculidae Clamator coromandus* | 22.3 | 9.08 | 9.37 | 161.6 | 25.67 | T | L | I | 77 |
| *Cuculidae Cuculus micropterus* | 20.03 | 7.85 | 9.13 | 197.1 | 19.44 | T | L | I | 119 |
| *Cuculidae Cuculus poliocephalus* | 14.12 | 6.1 | 6.03 | 147.6 | 15.89 | T | L | I | 47.5 |
| *Cuculidae Cuculus saturatus* | 17.3 | 7.27 | 8.24 | 185.9 | 17.29 | T | L | I | 90.6 |
| *Cuculidae Hierococcyx nisicolor* | 15.5 | 9.15 | 8 | 175 | 22.5 | M | L | I | 81.1 |
| *Cuculidae Hierococcyx sparverioides* | 18.87 | 8.48 | 8.31 | 221.8 | 26.69 | T | L | I | 151 |
| *Cuculidae Phaenicophaeus tristis* | 20.01 | 8.75 | 11.58 | 161 | 37.59 | B | B | I | 139 |
| *Cuculidae Surniculus lugubris* | 17.29 | 7.02 | 7.49 | 146 | 14.87 | T | L | I | 35 |
| *Dicaeidae Dicaeum cruentatum* | 7.4 | 2.88 | 2.9 | 49.8 | 12.23 | T | F | F | 6.2 |
| *Dicaeidae Dicaeum ignipectus* | 6.7 | 3.42 | 2.87 | 48.5 | 12.35 | T | F | F | 5.9 |
| *Dicaeidae Dicaeum melanozanthum* | 6 | 3.8 | 3.65 | 73.5 | 16 | T | F | F | 12.5 |
| *Dicruridae Dicrurus aeneus* | 14.01 | 8.14 | 6.13 | 126.1 | 15.38 | T | A | I | 23.1 |
| *Dicruridae Dicrurus annectens* | 15.35 | 9.89 | 9.28 | 145 | 20.94 | T | A | I | 44 |
| *Dicruridae Dicrurus hottentottus* | 25.64 | 10.58 | 11.16 | 152.6 | 25.28 | T | Fl | N | 79.2 |
| *Dicruridae Dicrurus leucophaeus* | 18.21 | 9.09 | 9.1 | 142.5 | 18.62 | T | A | I | 37.6 |
| *Dicruridae Dicrurus paradiseus* | 25.32 | 11.1 | 12.84 | 170 | 29.48 | T | A | I | 71 |
| *Dicruridae Dicrurus remifer* | 16.4 | 8.42 | 8.81 | 143 | 20.51 | T | A | I | 43.1 |
| *Elachuridae Elachura formosa* | 7.83 | 2.53 | 3.2 | 48 | 20.03 | B | L | I | 11 |
| *Eurylaimidae Psarisomus dalhousiae* | 15.9 | 12.93 | 9.5 | 101 | 27.7 | T | A | I | 67 |
| *Fringillidae Carpodacus sipahi* | 12.55 | 9.84 | 11.73 | 101.5 | 21.41 | M | F | S | 39.5 |
| *Fringillidae Carpodacus subhimachalus* | 9.06 | 9.17 | 9.81 | 95.5 | 22.33 | B | F | S | 47 |
| *Fringillidae Chloris spinoides* | 9.62 | 7.06 | 7.82 | 76.7 | 15.69 | B | F | S | 18.6 |
| *Fringillidae Leucosticte nemoricola* | 9.33 | 6.6 | 6.86 | 101 | 20.42 | G | G | S | 22.7 |
| *Fringillidae Mycerobas melanozanthos* | 19.24 | 17.06 | 20.23 | 132.5 | 25.32 | T | F | S | 62 |
| *Fringillidae Pyrrhoplectes epauletta* | 8.83 | 6.65 | 7.1 | 78 | 20 | B | F | S | 19 |
| *Fringillidae Pyrrhula erythaca* | 8.6 | 8.35 | 8.65 | 83 | 18.7 | B | F | F | 19 |
| *Fringillidae Pyrrhula nipalensis* | 9.71 | 10.13 | 9.85 | 88 | 17.07 | T | F | F | 24.6 |
| *Irenidae Irena puella* | 18.36 | 8.22 | 8.72 | 136.9 | 19.93 | T | F | F | 64.9 |
| *Leiotrichidae Actinodura egertoni* | 9.1 | 4.1 | 4.85 | 82.5 | 30.3 | M | L | O | 36 |
| *Leiotrichidae Alcippe nipalensis* | 7.26 | 3.08 | 3.81 | 59.1 | 23 | M | L | I | 15.8 |
| *Leiotrichidae Chrysominla strigula* | 7 | 3.45 | 3.96 | 66.2 | 26.1 | M | L | I | 19.2 |
| *Leiotrichidae Cutia nipalensis* | 12.15 | 4.7 | 6 | 94.8 | 34.95 | M | B | I | 47.5 |
| *Leiotrichidae Garrulax albogularis* | 15.76 | 6.27 | 7.29 | 132.3 | 44.45 | G | G | I | 98.5 |
| *Leiotrichidae Garrulax caerulatus* | 16.33 | 5.98 | 6.95 | 109.5 | 41.35 | B | F | F | 89 |
| *Leiotrichidae Garrulax leucolophus* | 16.8 | 5.76 | 7.82 | 136.5 | 47.64 | G | G | O | 123.5 |
| *Leiotrichidae Garrulax monileger* | 18.42 | 6.44 | 7.54 | 129.5 | 41.79 | G | G | I | 84 |
| *Leiotrichidae Garrulax ocellatus* | 19.55 | 6.29 | 7.7 | 130.5 | 47.68 | G | G | F | 120 |
| *Leiotrichidae Garrulax pectoralis* | 23.3 | 7.58 | 9.31 | 142.5 | 49.35 | G | G | I | 145.5 |
| *Leiotrichidae Grammatoptila striata* | 17.3 | 7.79 | 10.8 | 151 | 46.32 | M | L | I | 110.5 |
| *Leiotrichidae Heterophasia picaoides* | 15 | 4.53 | 4.87 | 118.5 | 28.05 | T | L | I | 42.4 |
| *Leiotrichidae Heterophasia pulchella* | 15.83 | 5.1 | 5.2 | 107 | 31.47 | T | L | I | 41 |
| *Leiotrichidae Leioptila annectens* | 8.8 | 3.65 | 4 | 80.5 | 25.7 | T | M | I | 24.5 |
| *Leiotrichidae Leiothrix argentauris* | 9.37 | 4.11 | 5.03 | 76 | 25.73 | B | L | I | 28.4 |

| | | | | | | | | | |
|---|---|---|---|---|---|---|---|---|---|
| *Leiotrichidae Liocichla phoenicea* | 9.8 | 4.2 | 5.68 | 87 | 35.3 | B | L | O | 49 |
| *Leiotrichidae Minla ignotincta* | 6.79 | 2.86 | 3.07 | 63.7 | 19.8 | M | L | I | 14.3 |
| *Leiotrichidae Sibia waldeni* | 10.68 | 4.28 | 5.1 | 95.5 | 33.85 | M | L | O | 47.5 |
| *Leiotrichidae Siva cyanouroptera* | 7.84 | 3.25 | 3.8 | 63.2 | 22.85 | M | L | I | 17 |
| *Leiotrichidae Trochalopteron affine* | 13.01 | 4.92 | 5.81 | 105 | 39.07 | G | G | F | 73 |
| *Leiotrichidae Trochalopteron erythrocephalum* | 13.91 | 4.79 | 5.83 | 103 | 40.23 | B | L | I | 71.7 |
| *Leiotrichidae Trochalopteron imbricatum* | 10.2 | 4.58 | 5.63 | 81 | 29.05 | G | G | O | 40.7 |
| *Leiotrichidae Trochalopteron squamatum* | 13.52 | 5.62 | 7.61 | 103 | 41.43 | G | G | O | 84 |
| *Leiotrichidae Trochalopteron subunicolor* | 9.89 | 4.5 | 6.15 | 94.8 | 37.14 | G | G | I | 66 |
| *Megalaimidae Psilopogon lineatus* | 25.89 | 11.45 | 14.28 | 128.7 | 34.76 | T | F | F | 149 |
| *Megalaimidae Psilopogon virens* | 32.96 | 13.7 | 17.06 | 140.3 | 33.78 | T | F | F | 202 |
| *Meropidae Nyctyornis athertoni* | 37.05 | 9.45 | 10.56 | 131.6 | 17.83 | T | A | I | 92 |
| *Monarchidae Hypothymis azurea* | 9.38 | 5.32 | 3.91 | 72 | 16.13 | M | A | I | 11.1 |
| *Monarchidae Terpsiphone paradisi* | 13.42 | 7.05 | 5.07 | 92.5 | 16.49 | M | A | I | 19.3 |
| *Muscicapidae Anthipes monileger* | 7.33 | 3.7 | 2.9 | 60.5 | 21.45 | B | A | I | 11 |
| *Muscicapidae Brachypteryx leucophris* | 7.65 | 2.96 | 3.1 | 57.8 | 27.85 | G | G | I | 15.8 |
| *Muscicapidae Brachypteryx montana* | 8.28 | 3.15 | 3.25 | 66.9 | 32.44 | G | G | I | 17.7 |
| *Muscicapidae Cinclidium frontale* | 9.32 | 3.15 | 3.82 | 93.5 | 27.9 | G | G | I | 25.4 |
| *Muscicapidae Copsychus malabaricus* | 11.82 | 4.2 | 4.95 | 92.5 | 26.14 | G | G | I | 29.4 |
| *Muscicapidae Cyornis poliogenys* | 8.9 | 4.71 | 3.76 | 73.9 | 19.46 | B | A | I | 15.7 |
| *Muscicapidae Cyornis unicolor* | 9.58 | 4.99 | 3.97 | 81.2 | 17.56 | M | A | I | 21 |
| *Muscicapidae Enicurus immaculatus* | 12.59 | 4.18 | 3.88 | 90 | 25.6 | R | G | I | 25.5 |
| *Muscicapidae Enicurus maculatus* | 15.1 | 4.48 | 4.55 | 106.5 | 29.47 | R | G | I | 41 |
| *Muscicapidae Enicurus schistaceus* | 13.14 | 4.55 | 4.06 | 100 | 29.11 | R | G | I | 31 |
| *Muscicapidae Eumyias thalassinus* | 6.74 | 4.52 | 3.22 | 83.4 | 15.85 | T | A | I | 17.7 |
| *Muscicapidae Ficedula hodgsoni* | 6.09 | 2.43 | 2.35 | 53.1 | 16.1 | B | A | I | 5 |
| *Muscicapidae Ficedula sapphira* | 6.28 | 3.35 | 2.55 | 62 | 16.4 | T | A | I | 7.8 |
| *Muscicapidae Ficedula strophiata* | 6.41 | 3.28 | 3.01 | 74.1 | 20 | M | A | I | 13 |
| *Muscicapidae Ficedula tricolor* | 6.07 | 3.08 | 2.56 | 60.4 | 19.25 | B | A | I | 8.3 |
| *Muscicapidae Ficedula westermanni* | 6.3 | 3.31 | 2.78 | 58.1 | 15.48 | T | A | I | 7.8 |
| *Muscicapidae Monticola rufiventris* | 14.37 | 5.56 | 6.09 | 122.7 | 28.03 | T | G | I | 53.2 |
| *Muscicapidae Muscicapa ferruginea* | 6.09 | 4.48 | 2.96 | 69.2 | 14.5 | M | A | I | 12 |
| *Muscicapidae Muscicapa sibirica* | 5.4 | 3.59 | 2.51 | 73.1 | 11.75 | M | A | I | 13.2 |
| *Muscicapidae Myiomela leucura* | 10 | 3.43 | 4.26 | 95.6 | 27.88 | G | G | I | 25 |
| *Muscicapidae Myophonus caeruleus* | 18.85 | 7.05 | 9.78 | 172 | 52.13 | R | G | I | 158 |
| *Muscicapidae Niltava grandis* | 10.93 | 5.33 | 5.03 | 104.4 | 25.2 | M | A | I | 30.3 |
| *Muscicapidae Niltava macgrigoriae* | 6.37 | 3.57 | 3.11 | 64.3 | 16.7 | B | A | I | 11.9 |
| *Muscicapidae Niltava sundara* | 8.42 | 4.35 | 3.86 | 80.7 | 21.6 | B | A | I | 21.1 |
| *Muscicapidae Phoenicurus frontalis* | 7.27 | 3.01 | 2.94 | 86.2 | 24.4 | G | G | I | 16 |
| *Nectariniidae Aethopyga gouldiae* | 12.2 | 3.2 | 2.3 | 56 | 14 | T | Fl | N | 6.1 |
| *Nectariniidae Aethopyga nipalensis* | 16.65 | 3 | 3.1 | 54.5 | 16.1 | T | Fl | N | 7 |
| *Nectariniidae Aethopyga saturata* | 15.95 | 3 | 3.1 | 52 | 14.55 | T | Fl | N | 5.3 |
| *Nectariniidae Aethopyga siparaja* | 15.95 | 3.7 | 2.85 | 62 | 14.7 | T | Fl | N | 6.7 |
| *Nectariniidae Arachnothera magna* | 35.95 | 6.1 | 5.3 | 94.3 | 21.7 | M | Fl | N | 30.7 |
| *Oriolidae Oriolus traillii* | 22.02 | 7.3 | 7.99 | 148.7 | 24.27 | T | F | F | 74 |
| *Oriolidae Oriolus xanthornus* | 22.07 | 8.01 | 8.23 | 140.9 | 26.11 | T | F | F | 56.3 |
| *Paridae Machlolophus spilonotus* | 8.42 | 4.57 | 5.02 | 77.2 | 19.9 | T | L | I | 18.8 |
| *Paridae Melanochlora sultanea* | 11.38 | 5.42 | 14.46 | 108.6 | 24.4 | T | L | I | 37.6 |
| *Paridae Sylviparus modestus* | 5.05 | 2.65 | 2.84 | 59.5 | 17.55 | T | L | I | 8.2 |
| *Pellorneidae Gampsorhynchus rufulus* | 11.34 | 5.15 | 6.34 | 98 | 28.01 | B | L | I | 37 |
| *Pellorneidae Malacocincla abbotti* | 12.68 | 4 | 5.8 | 78 | 26.3 | B | L | I | 29.3 |
| *Pellorneidae Napothera epilepidota* | 10.94 | 3.09 | 3.55 | 54.9 | 22.24 | G | G | I | 16 |
| *Pellorneidae Pellorneum ruficeps* | 10.9 | 3.4 | 4.85 | 71 | 27.3 | G | G | I | 26 |
| *Pellorneidae Rimator malacoptilus* | 17.65 | 2.9 | 3.35 | 60.5 | 24.5 | G | G | I | 19.5 |
| *Pellorneidae Schoeniparus castaneceps* | 6.6 | 2.28 | 2.75 | 57.5 | 22.2 | B | L | I | 12.5 |
| *Phylloscopidae Phylloscopus cantator* | 7.96 | 3 | 2.53 | 52.8 | 16.34 | T | L | I | 6 |

| | | | | | | | | | |
|---|---|---|---|---|---|---|---|---|---|
| *Phylloscopidae Phylloscopus castaniceps* | 5.1 | 2.35 | 2.38 | 51.2 | 16.3 | T | L | I | 5.3 |
| *Phylloscopidae Phylloscopus chloronotus* | 5.44 | 2.56 | 1.95 | 54.5 | 16.59 | T | L | I | 5.1 |
| *Phylloscopidae Phylloscopus intermedius* | 7.38 | 3.1 | 2.84 | 56.5 | 16.93 | B | A | I | 8 |
| *Phylloscopidae Phylloscopus maculipennis* | 5.31 | 2.46 | 1.92 | 50.8 | 17.86 | T | L | I | 5.1 |
| *Phylloscopidae Phylloscopus magnirostris* | 8.1 | 3.81 | 3.1 | 69.7 | 20.27 | B | L | I | 11.6 |
| *Phylloscopidae Phylloscopus poliogenys* | 6.83 | 3.13 | 2.57 | 51.2 | 17.32 | B | A | I | 6.3 |
| *Phylloscopidae Phylloscopus pulcher* | 6.55 | 2.56 | 2.37 | 59.3 | 18.97 | M | L | I | 6.8 |
| *Phylloscopidae Phylloscopus reguloides* | 7.4 | 3.21 | 2.62 | 58.5 | 17.79 | T | L | I | 7.8 |
| *Phylloscopidae Phylloscopus whistleri* | 7.35 | 3.15 | 2.76 | 58.3 | 19.08 | B | A | I | 7.3 |
| *Phylloscopidae Phylloscopus xanthoschistos* | 7.26 | 2.8 | 2.61 | 56.9 | 19.16 | T | L | I | 7 |
| *Picidae Blythipicus pyrrhotis* | 36.45 | 9.35 | 11.3 | 148.5 | 26.5 | G | G | I | 132 |
| *Picidae Chrysocolaptes lucidus* | 41.2 | 11.7 | 12.7 | 179 | 30.5 | T | B | I | 142 |
| *Picidae Chrysophlegma flavinucha* | 29 | 10.15 | 10.1 | 171 | 28.9 | M | B | I | 177.3 |
| *Picidae Dendrocopos cathpharius* | 14.2 | 4.95 | 5.35 | 100.8 | 17.2 | T | B | I | 31 |
| *Picidae Dendrocopos darjellensis* | 28.13 | 9.06 | 7.4 | 133 | 23.75 | T | B | I | 71 |
| *Picidae Dendrocopos hyperythrus* | 24.86 | 5.93 | 6.25 | 124 | 20.75 | T | B | I | 46.5 |
| *Picidae Dendrocopos macei* | 20.4 | 6.3 | 6.4 | 108.8 | 18.58 | M | B | I | 40.6 |
| *Picidae Gecinulus grantia* | 19.3 | 8.6 | 8.4 | 130 | 25.3 | M | B | I | 76.5 |
| *Picidae Micropternus brachyurus* | 18.2 | 7.25 | 7.95 | 131.5 | 21.3 | T | B | I | 69.5 |
| *Picidae Mulleripicus pulverulentus* | 47 | 13.25 | 13.75 | 240.5 | 40.2 | T | B | I | 461.5 |
| *Picidae Picumnus innominatus* | 10.96 | 4.46 | 4.57 | 59 | 11.49 | M | B | I | 10.2 |
| *Picidae Picus canus* | 28.3 | 9.65 | 9.35 | 141.5 | 28.95 | M | B | I | 137 |
| *Picidae Picus chlorolophus* | 22.55 | 7.6 | 7.6 | 139 | 23.15 | M | B | I | 67 |
| *Picidae Sasia ochracea* | 11 | 3.95 | 4.7 | 54 | 13.35 | M | B | I | 9.7 |
| *Pittidae Hydrornis nipalensis* | 19.65 | 7.7 | 10.15 | 123 | 54 | G | G | I | 124 |
| *Pittidae Pitta sordida* | 15.05 | 6.5 | 8.35 | 111 | 40.85 | G | G | I | 64.5 |
| *Pnoepygidae Pnoepyga albiventer* | 8.01 | 3.3 | 3.12 | 59.4 | 24.19 | G | G | I | 20.9 |
| *Pnoepygidae Pnoepyga pusilla* | 8.26 | 3.2 | 2.67 | 48.3 | 19.24 | G | G | I | 12 |
| *Podargidae Batrachostomus hodgsoni* | 12.27 | 14.74 | 5.74 | 115.5 | 15.56 | M | G | I | 51 |
| *Psittacidae Psittacula alexandri* | 26.04 | 15.61 | 25.54 | 169 | 15.89 | T | F | F | 150.5 |
| *Pycnonotidae Alophoixus flaveolus* | 11.28 | 4.75 | 6.59 | 103.5 | 22.3 | M | F | F | 45.7 |
| *Pycnonotidae Hemixos flavala* | 12.75 | 5.21 | 5.39 | 97.3 | 19.05 | T | F | F | 32.5 |
| *Pycnonotidae Hypsipetes leucocephalus* | 17.04 | 5.83 | 6.12 | 123.8 | 19.42 | T | F | F | 52.5 |
| *Pycnonotidae Ixos mcclellandii* | 18.11 | 5.94 | 5.7 | 107.5 | 17.85 | T | F | F | 34 |
| *Pycnonotidae Pycnonotus flaviventris* | 8.8 | 4.21 | 4.49 | 91.3 | 16.93 | M | F | F | 27.8 |
| *Pycnonotidae Pycnonotus striatus* | 10.06 | 4.89 | 5.34 | 109.8 | 19.72 | T | F | F | 52.5 |
| *Ramphastidae Megalaima asiatica* | 17.27 | 9.48 | 11 | 103.5 | 26.75 | T | F | F | 90.5 |
| *Ramphastidae Megalaima australis* | 14.3 | 7.6 | 8.14 | 86.3 | 19.51 | T | F | F | 33.3 |
| *Ramphastidae Megalaima franklinii* | 17.95 | 10.2 | 11.18 | 103.9 | 26.87 | T | F | F | 63.5 |
| *Rhipiduridae Rhipidura albicollis* | 7.48 | 4.11 | 2.9 | 78 | 20.44 | B | A | I | 12.9 |
| *Scotocercidae Abroscopus albogularis* | 5.48 | 2.5 | 2.1 | 43.5 | 15.6 | M | L | I | 5 |
| *Scotocercidae Abroscopus schisticeps* | 5.5 | 2.88 | 2.25 | 49.5 | 16 | M | L | I | 4.7 |
| *Scotocercidae Abroscopus superciliaris* | 6.5 | 2.7 | 2.7 | 50.8 | 17.15 | M | L | I | 6.5 |
| *Scotocercidae Cettia brunnifrons* | 5.26 | 1.98 | 2.03 | 45.4 | 18.32 | B | L | I | 7.5 |
| *Scotocercidae Cettia castaneocoronata* | 6.38 | 2.72 | 2.28 | 50.5 | 22.02 | G | G | I | 8.8 |
| *Scotocercidae Horornis brunnescens* | 6.47 | 2.23 | 2.18 | 52.7 | 21.85 | B | L | I | 6 |
| *Scotocercidae Horornis fortipes* | 5.9 | 2.17 | 2.43 | 54.3 | 22.5 | B | L | I | 10.4 |
| *Scotocercidae Phyllergates cucullatus* | 9.52 | 2.75 | 2.27 | 48.3 | 20.07 | B | M | I | 5.9 |
| *Scotocercidae Tesia cyaniventer* | 7.92 | 3.81 | 2.76 | 49.6 | 24.67 | G | G | I | 9.7 |
| *Scotocercidae Tesia olivea* | 7.32 | 3.15 | 2.72 | 45.1 | 21.63 | G | G | I | 7 |
| *Scotocercidae Tickellia hodgsoni* | 6.65 | 2.99 | 2.18 | 48 | 23.07 | B | A | I | 4.5 |
| *Sittidae Sitta cinnamoventris* | 14.03 | 4.63 | 4.8 | 81.8 | 19.88 | T | B | I | 19.6 |
| *Sittidae Sitta formosa* | 13.8 | 5 | 5.1 | 106 | 21.9 | T | B | I | 24.7 |
| *Sittidae Sitta himalayensis* | 10.35 | 5.18 | 3.48 | 73 | 17.75 | T | B | I | 14.5 |
| *Stenostiridae Chelidorhynx hypoxanthus* | 4.05 | 2.58 | 1.75 | 57.8 | 15.58 | T | A | I | 5.5 |
| *Stenostiridae Culicicapa ceylonensis* | 6.2 | 3.53 | 2.49 | 62.5 | 13.8 | M | A | I | 7.7 |

| | | | | | | | | | |
|---|---|---|---|---|---|---|---|---|---|
| *Strigidae Glaucidium brodiei* | 9.85 | 5.21 | 8.99 | 91 | 19.8 | M | A | I | 58 |
| *Strigidae Glaucidium cuculoides* | 13.83 | 10.02 | 12.49 | 150.5 | 29.58 | M | G | I | 162.5 |
| *Sturnidae Acridotheres tristis* | 17.46 | 7.77 | 9.23 | 144 | 41.94 | G | G | O | 116.6 |
| *Sturnidae Gracula religiosa* | 20 | 10.39 | 11.75 | 168.5 | 36.19 | T | F | O | 192 |
| *Sylviidae Chleuasicus atrosuperciliaris* | 6.85 | 6.28 | 8.55 | 71.5 | 25.85 | B | L | I | 16.8 |
| *Sylviidae Cholornis unicolor* | 10.85 | 6.89 | 10.11 | 90.3 | 30.75 | B | L | S | 34 |
| *Sylviidae Fulvetta ludlowi* | 5.75 | 2.43 | 3.33 | 56.5 | 23.5 | B | L | I | 12.3 |
| *Sylviidae Lioparus chrysotis* | 4.87 | 2.65 | 3.03 | 51.7 | 20.63 | B | L | I | 11.8 |
| *Sylviidae Myzornis pyrrhoura* | 9.75 | 3.12 | 2.59 | 62.3 | 24.6 | B | I | I | 11.9 |
| *Sylviidae Paradoxornis gularis* | 9.63 | 6.06 | 9.22 | 87 | 26.46 | B | L | S | 29 |
| *Sylviidae Psittiparus ruficeps* | 10.3 | 7.1 | 12.2 | 85 | 25.4 | B | L | I | 32 |
| *Sylviidae Suthora nipalensis* | 4.54 | 4.19 | 5.78 | 50.5 | 17.43 | B | L | I | 5.5 |
| *Timaliidae Cyanoderma chrysaeum* | 7.6 | 2.8 | 3.35 | 53 | 20.6 | B | L | I | 9 |
| *Timaliidae Cyanoderma ruficeps* | 8.95 | 2.8 | 3.3 | 55.3 | 20.6 | B | L | I | 10.3 |
| *Timaliidae Macronous gularis* | 8.68 | 3.1 | 3.5 | 60 | 19.3 | B | L | I | 11.5 |
| *Timaliidae Pomatorhinus ferruginosus* | 18.8 | 3.9 | 6.4 | 89.5 | 33.7 | G | G | I | 40 |
| *Timaliidae Pomatorhinus ruficollis* | 15.9 | 3.63 | 6 | 82.5 | 28.55 | G | G | I | 31.7 |
| *Timaliidae Pomatorhinus schisticeps* | 20.93 | 4 | 6.8 | 98.5 | 34.05 | G | G | I | 43 |
| *Timaliidae Pomatorhinus superciliaris* | 43.15 | 2.7 | 5 | 77 | 31.35 | G | G | I | 28 |
| *Timaliidae Spelaeornis caudatus* | 6.46 | 2.75 | 3.24 | 48.3 | 19.2 | B | L | I | 11 |
| *Timaliidae Spelaeornis troglodytoides* | 5.93 | 2.3 | 3 | 51 | 22.45 | B | L | I | 11 |
| *Timaliidae Sphenocichla humei* | 17.5 | 4.5 | 6.7 | 73 | 28.1 | B | L | I | 34.1 |
| *Timaliidae Stachyris nigriceps* | 9.93 | 3.43 | 4.65 | 58 | 23.05 | B | L | I | 15.8 |
| *Trogonidae Harpactes erythrocephalus* | 12.4 | 8.66 | 10.53 | 153.5 | 17.08 | M | A | I | 80.3 |
| *Trogonidae Harpactes wardi* | 10.85 | 7.95 | 10.75 | 161 | 16.7 | M | A | I | 119 |
| *Turdidae Cochoa purpurea* | 10 | 6 | 5.7 | 140 | 27.8 | T | L | O | 103 |
| *Turdidae Cochoa viridis* | 11.4 | 6.8 | 5.9 | 142 | 30.2 | T | L | O | 106.8 |
| *Vangidae Hemipus picatus* | 9.68 | 4.48 | 3.57 | 64.4 | 12.7 | T | A | I | 9 |
| *Vangidae Tephrodornis virgatus* | 16.13 | 6.98 | 7.43 | 118.8 | 20.66 | T | L | I | 37.8 |
| *Vireonidae Erpornis zantholeuca* | 9.93 | 3.02 | 3.97 | 67 | 16.1 | T | L | I | 11.8 |
| *Vireonidae Pteruthius flaviscapis* | 9.68 | 5.27 | 6.24 | 83 | 26.98 | T | L | I | 39 |
| *Vireonidae Pteruthius melanotis* | 5.34 | 3.25 | 3.97 | 60.3 | 19.57 | M | L | I | 13.3 |
| *Vireonidae Pteruthius rufiventer* | 9.8 | 4.88 | 6.39 | 88 | 28.53 | M | L | I | 44.5 |
| *Vireonidae Pteruthius xanthochlorus* | 5.05 | 3.36 | 3.69 | 61.5 | 20.6 | M | L | I | 14.3 |
| *Zosteropidae Yuhina bakeri* | 6.5 | 3.4 | 4.03 | 67 | 21.79 | M | L | I | 17.5 |
| *Zosteropidae Yuhina castaniceps* | 5.62 | 2.95 | 3.08 | 61.3 | 17.15 | M | L | I | 13.5 |
| *Zosteropidae Yuhina flavicollis* | 6.7 | 2.93 | 3.22 | 63.2 | 21.72 | M | L | I | 15.4 |
| *Zosteropidae Yuhina gularis* | 8.29 | 3.2 | 3.13 | 73.1 | 23.85 | T | L | I | 21 |
| *Zosteropidae Yuhina nigrimenta* | 6.74 | 2.63 | 2.52 | 56 | 16.4 | M | L | I | 9.5 |
| *Zosteropidae Yuhina occipitalis* | 8.16 | 2.75 | 2.95 | 61.5 | 20.09 | T | L | I | 13 |

The entire set of environmental data was obtained from CHELSA-CLIMATOLOGIES available at *http://chelsa-climate.org/bioclim/*. The following bioclimatic variables were downloaded:

- MAT – Mean Annual Temperature
- TMAX – Maximum Temperature of the warmest month
- TMIN – Minimum Temperature of the warmest month
- APPT – Annual mean precipitation
- CVPPT – Precipitation seasonality (Coefficient of Variation)

For the contemporary climate, a single averaged raster per variable was downloaded for 1979-2013. For assembling the paleoclimatic variables, three common Global Climate Models (GCMs) were used – Community Climate System Model 4 (CCSM4; Gent et al., 2011), Model for Interdisciplinary Research on Climate – Earth System Model (MIROC-ESM; Watanabe et al., 2011) and the Max Planck Institute – Earth System Model running in low resolution grid and paleo mode (MPI-ESM-P; Giorgetta et al., 2013). The Quaternary climatic stability was defined as the change in the contemporary and historic variable (Jansson, 2003) and was averaged across the three GCMs.

**Table S3.** Environmental variables of sampled communities. Sites = Details of the bird elevational transects/ hawkmoth light screens; Plot = elevational code, Lat = Latitude, Long = Longitude , EVI: enhanced vegetation index (productivity); MAT = mean annual temperature, TMAX = maximum temperature of the hottest month, TMIN = minimum temperature of the coldest month, APPT = mean annual precipitation; CVPPT = Coefficient of variation in precipitation, Quaternary Climatic stability = Difference between the environmental variables assembled using Last Glacial Maxima climatologies from CHELSA (http://chelsa-climate.org/bioclim/) and the contemporary climate.

| | | | | **Contemporary Climate (1980 - 2014)** | | | | | **Quaternary Climatic Stablility** | | | | |
|---|---|---|---|---|---|---|---|---|---|---|---|---|---|
| **Site** | Lat | Long | EVI | MAT | TMAX | TMIN | APPT | CVPPT | ΔMAT | ΔTMAX | ΔTMIN | ΔAPPT | ΔCVPPT |
| **E0200** | 92.6017 | 27.0375 | 0.4373 | 23.1 | 30.8 | 12.0 | 1855 | 83 | 9.6 | 5.2 | 15.7 | 30 | -6.7 |
| **E0500** | 92.4186 | 26.9956 | 0.3812 | 21.7 | 29.4 | 10.7 | 2397 | 87 | 9.8 | 5.5 | 16.0 | 613 | -1.7 |
| **E0700** | 92.4162 | 27.0071 | 0.3881 | 20.7 | 28.5 | 9.7 | 2429 | 87 | 10.1 | 5.8 | 16.25 | 533 | -2.4 |
| **E0900** | 92.4141 | 27.0199 | 0.3555 | 18.9 | 26.8 | 7.9 | 2559 | 86 | 10.4 | 6.2 | 16.6 | 364 | -5.1 |
| **E1100** | 92.4132 | 27.0243 | 0.3599 | 18.9 | 26.8 | 7.9 | 2559 | 86 | 10.4 | 6.2 | 16.6 | 364 | -5.1 |
| **E1300** | 92.4156 | 27.0372 | 0.4026 | 18 | 25.9 | 7.0 | 2616 | 85 | 10.6 | 6.3 | 16.7 | 501 | -3.7 |
| **E1500** | 92.4107 | 27.0594 | 0.3207 | 15.7 | 23.8 | 4.7 | 2726 | 85 | 10.9 | 6.9 | 17.25 | 299 | -6.8 |
| **E1700** | 92.4136 | 27.0657 | 0.3597 | 15.7 | 23.8 | 4.7 | 2726 | 85 | 10.9 | 6.9 | 17.25 | 299 | -6.8 |
| **E1900** | 92.4055 | 27.0665 | 0.4078 | 15 | 23.1 | 4.0 | 2732 | 85 | 11.2 | 7.1 | 17.45 | 189 | -6.3 |
| **E2100** | 92.4071 | 27.073 | 0.3509 | 13.1 | 21.3 | 2.1 | 2806 | 84 | 11.5 | 7.5 | 17.81 | 105 | -7.3 |
| **E2300** | 92.4056 | 27.0798 | 0.2981 | 12.3 | 20.5 | 1.3 | 2858 | 84 | 11.7 | 7.8 | 18.08 | 122 | -6.7 |
| **E2500** | 92.4369 | 27.1146 | 0.3142 | 11.1 | 19.4 | 0.1 | 2931 | 83 | 11.9 | 8.1 | 18.31 | 197.1 | -9.9 |
| **E2770** | 92.45 | 27.1312 | 0.2397 | 9.7 | 18 | -1.3 | 2795 | 82 | 12.4 | 8.4 | 18.72 | -96 | -12.7 |

**Appendix S2**

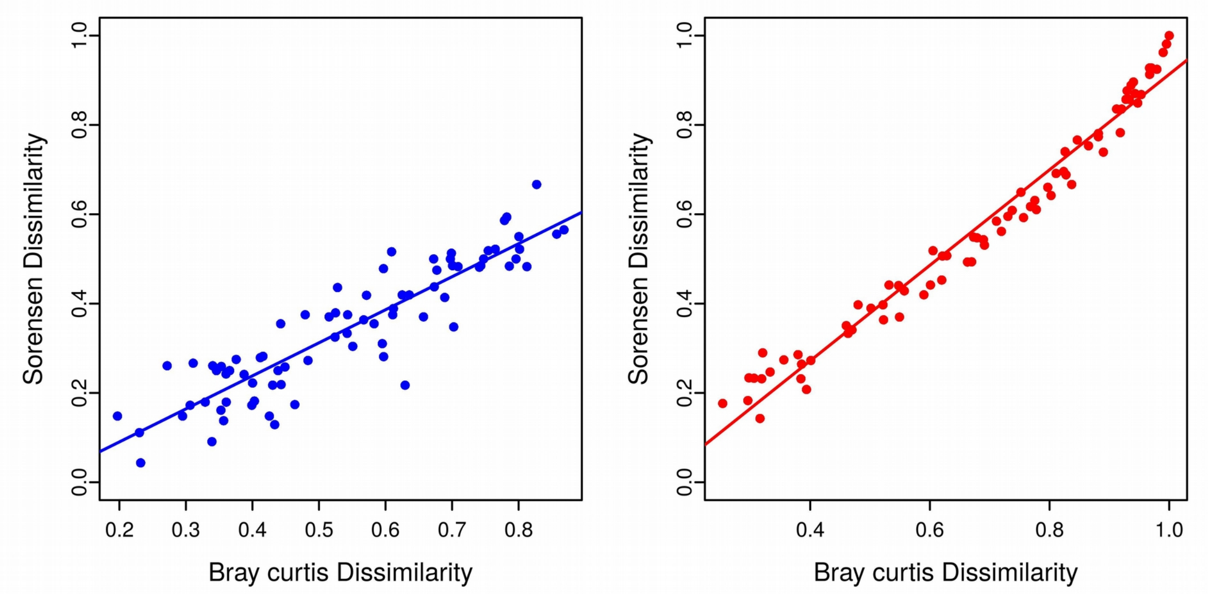


**Figure S4.** Ordinary least squares regression fits between the taxonomic dissimilarities calculated using the abundance based Bray curtis index and the abundance based SØrensen index. Plots are presented for hawkmoth (left; blue) and birds (right; red). The coefficients for the linear fits are listed in the table below.

| | **Intercept** | **Slope** | $r^2$ | **p.value** |
|---|---|---|---|---|
| **Hawkmoths** | -0.06 ± 0.02 | 0.74 ± 0.04 | 0.81 | < 0.005 |
| **Birds** | -0.16 ± 0.02 | 1.07 ± 0.02 | 0.97 | < 0.005 |

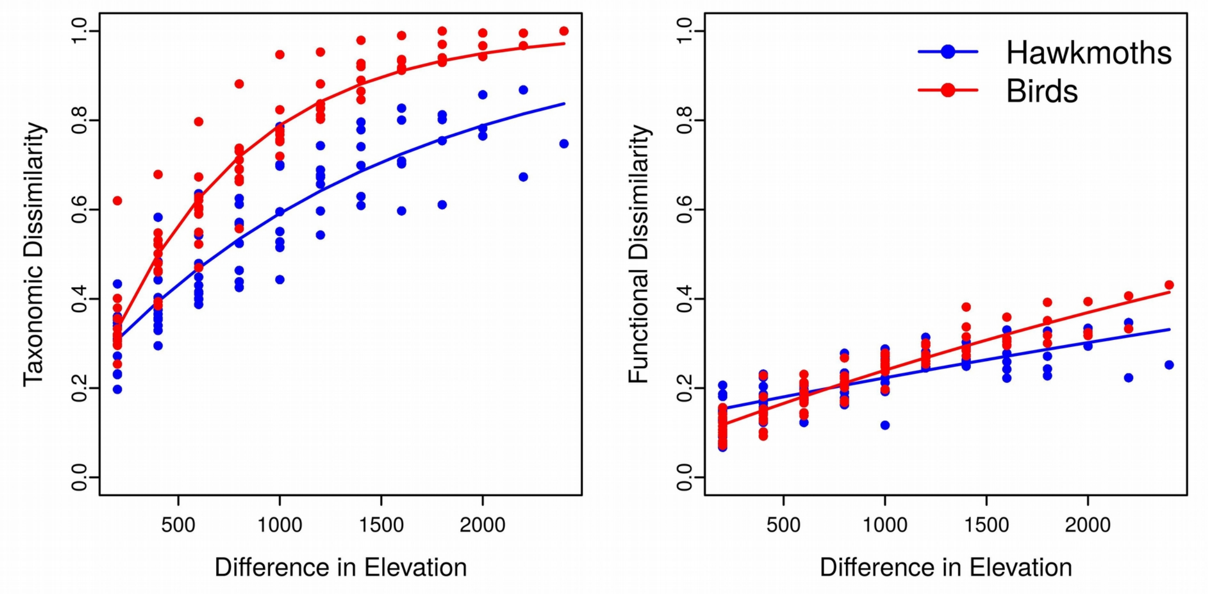


**Figure S5**. A comparison of slopes between the distance decay relationships for taxonomic (left) and functional (right) dissimilarities. The blue points and line correspond to hawkmoth dissimilarity and the red points and line represent bird data sets. The difference in slope was significant between the hawkmoths and birds dissimilarity (taxonomic - Fisher's z = 8.68, p.value < 0.001; functional - Fisher's z = 6.03, p.value < 0.001), indicating an overall higher rate of turnover for birds. The coefficients for the quadratic fits ($\beta \sim 1 - a * \exp(b * \text{distance})$ are listed in the table below

| | | **a** | **b** | **p.value** |
|---|---|---|---|---|
| **Taxonomic** | **Hawkmoths** | 0.79 ± 0.002 | $(-6.6 \pm 0.4) \times 10^{-4}$ | < 0.005 |
| | **Birds** | 0.89 ± 0.003 | $(-1.4 \pm 0.08) \times 10^{-3}$ | < 0.005 |
| **Functional** | **Hawkmoths** | 0.86 ± 0.009 | $(-1.1 \pm 0.1) \times 10^{-4}$ | < 0.005 |
| | **Birds** | 0.91 ± 0.07 | $(-1.9 \pm 0.08) \times 10^{-4}$ | < 0.005 |

**Table S4.** The various functions and R libraries associated with each analysis presented in the paper are listed below. All analyses were performed in R 3.4.4 on a Ubuntu (linux-gnu) 18.04.1 platform (R Core Development Team 2013).

| **Analysis** | **Function** | **Package** | **Reference** |
|---|---|---|---|
| Partitioning Bray-curtis taxonomic dissimilarity | *bray.part* | betapart 1.5.1 | Baselga et al., 2018 |
| Species x Species distance matrix based on traits (categorical and numeric) | *gowdis* | vegan 2.5-5 | Oksanen et al. 2007 |
| cluster analysis | *hclust* | base R | |
| Generation of functional tree | *as.phylo.hclust* | ape 5.2 | Paradis & Schliep 2018 |
| Partitioning SØrensen functional dissimilarities | *part.p.tree* | commEcol 1.7.0 | Melo 2017 |
| Mantel tests | *mantel.test* | base stats | |
| *GDM model fitting, variable importance and ispline extractions* | *gdm, gdm.varImp, isplineExtract* | *gdm 1.3.11* | *Manion et al., 2018* |
| *partial mantel tests* | *partial.mantel.test* | *ncf 1.2-8* | *Bjornstad & Cai, 2019* |

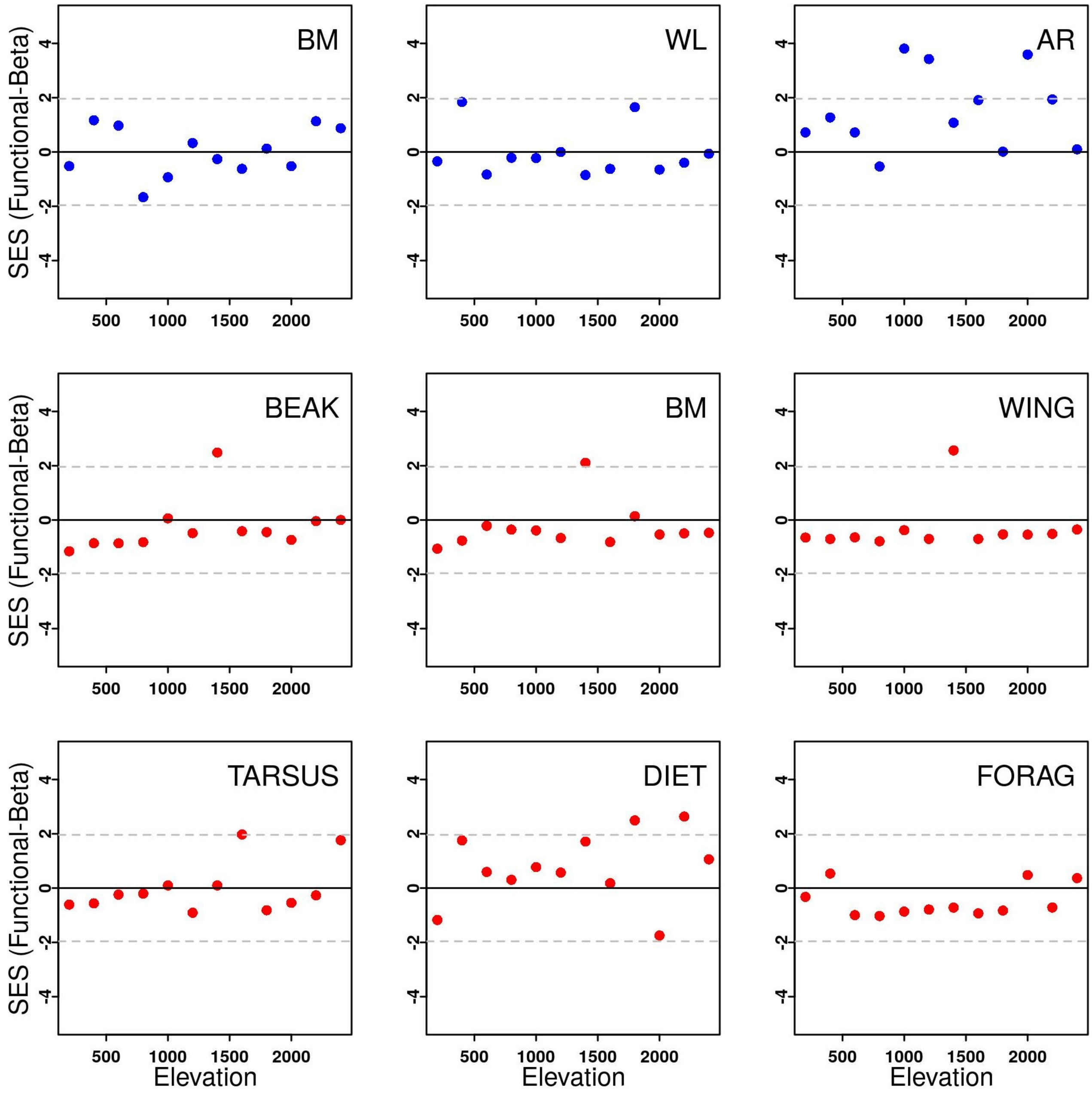


**Figure S6.** SES values for the deviation of observed functional beta-diversity using individual traits for hawkmoths (blue; top row) and birds (red; middle and bottom row). SES values greater than 1.96, or less than -1.96 (the dashed grey lines) indicate that the observed values of the respective communities are higher or lower than the values obtained under randomized assembly (see main text for details). Traits = BM = Body mass, WL = Wing loading, AR = Wing aspect ratio, BEAK = beak dimension, WING = wing length, TARSUS = length from the base of the tarsus to the first scute, DIET = primary diet of the species during breeding season, FORAG = Foraging mode. See Table S1.2 above for details on individual traits.

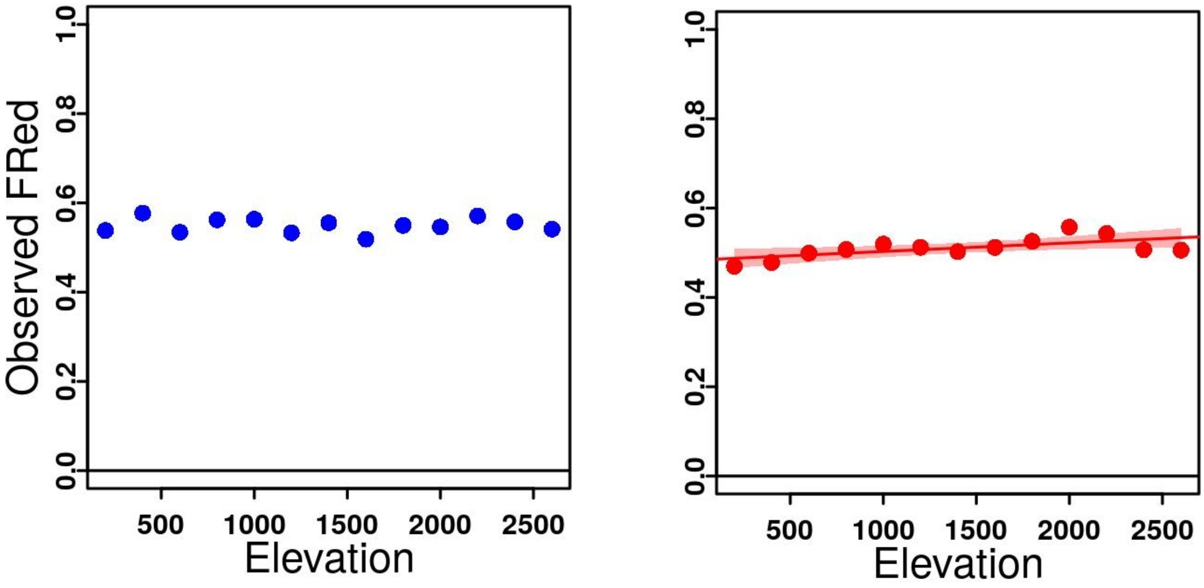


**Figure S7.**Observed functional redundancy values for hawkmoths (left; blue) and birds (right; red) across the elevational gradient

**Table S5.** Coefficients from the partial mantel tests for taxonomic and functional dissimilarities of hawkmoths and birds; independent matrices used were Env. Mat (environmental distance matrix) and Geo.Mat (Geographic distance matrix).

| **Model** | | **Env. Mat** | | **Geo. Mat** | |
|---|---|---|---|---|---|
| | | **Mantel's R** | **p.val** | **Mantel's R** | **p.val** |
| **birds** | Taxonomic beta ~ env. mat. + geo. mat. | 0.20 | 0.06 | 0.61 | <0.005 |
| | Functional beta ~ env. mat. + geo. mat. | 0.14 | 0.17 | 0.66 | < 0.005 |
| **hawkmoths** | Taxonomic beta ~ env. mat. + geo. mat. | 0.11 | 0.18 | 0.48 | < 0.005 |
| | Functional beta ~ env. mat. + geo. mat. | 0.12 | 0.18 | 0.29 | < 0.05 |

**Table S6.** Coefficients from the Multiple Regression on distance matrices for taxonomic and functional dissimilarities of hawkmoths and birds; independent matrices used were Env. Mat (environmental distance matrix) and Geo.Mat (Geographic distance matrix).

| **Model** | | **Env. Mat** | | **Geo. Mat** | | **r.sqrd** | **p.val** |
|---|---|---|---|---|---|---|---|
| | | **Coef** | **p.val** | **Coef** | **p.val** | | |
| **birds** | Taxonomic beta ~ env. mat. + geo. mat. | $3.73 \times 10^{-2}$ | 0.10 | $2.46 \times 10^{-5}$ | < 0.05 | 0.60 | < 0.05 |
| | Functional beta ~ env. mat. + geo. mat. | $9.44 \times 10^{-3}$ | 0.34 | $1.05 \times 10^{-5}$ | < 0.01 | 0.63 | < 0.05 |
| **hawkmoths** | Taxonomic beta ~ env. mat. + geo. mat. | $1.80 \times 10^{-2}$ | 0.47 | $1.66 \times 10^{-5}$ | < 0.05 | 0.41 | < 0.05 |
| | Functional beta ~ env. mat. + geo. mat. | $8.53 \times 10^{-3}$ | 0.37 | $3.74 \times 10^{-6}$ | 0.05 | 0.21 | < 0.05 |
